# Simultaneous 2- and 3-photon multiplane imaging across cortical layers in freely moving mice

**DOI:** 10.1101/2025.08.01.668113

**Authors:** Alexandr Klioutchnikov, Damian J. Wallace, Caleb Berdahl, Adam Sugi, Juergen Sawinski, Jason N. D. Kerr

**Author notes:** Correspondence to: Jason Kerr or Alexandr Klioutchnikov.

## Abstract

Head-mounted multiphoton microscopes enable imaging of activity from neuronal populations spread throughout the cortical layers in freely moving mice, but so far have been restricted to recording from one cortical layer at a time. Here, combining 2- and 3-photon based excitation delivered through multiple fibers, we built a head-mounted multiplane microscope enabling near simultaneous imaging (8ns between planes) of neuronal activity from five vertically separated planes, spread across multiple cortical layers. Both excitation pathways had remote focusing mechanisms for fine axial adjustments enabling activity recordings from the same neuronal populations over weeks in freely behaving mice. The lightweight microscope utilized an onboard, 2-channel detection system designed to enable activity recordings from neuronal populations spread across visual-cortex layers in both lit and dark conditions as well as imaging activity across posterior parietal cortex layers during complex gap-crossing behaviors. We show that during gap-crossing tasks, layer 5 and 2/3 neuronal subpopulations in posterior parietal cortex have differential pattern sequences during free decision making.

**Editorial summary:** A light-weight head-mounted multiplane microscope allows simultaneous imaging from >1800 neurons spread across the cortical layers in freely moving mice performing complex behavioral tasks, sampled over weeks.

## Main

To understand the cortical transformation of sensory-derived inputs, theoretical frameworks^1–3^ of cortical function have assigned a clear functional role in transforming sensory-derived input to each cortical layer based on the distinct physiological^4–6^ and anatomical^7^ properties of the layer’s neuronal inhabitants^8^. Over the past decade, high-resolution tools have been developed to measure activity from large numbers of neurons spread throughout the various cortical layers in the freely behaving animal with the aim of exploring the link between complex behavior and the underlying neuronal activity. Large scale extracellular electrical recordings have shown differences in cortical modulation^9^ and have unparalleled temporal resolution across vast numbers of recording sites^10^, but given their inability to unambiguously identify cell types^11^ or spatial position^12^, relating the recordings to connectomic reconstruction of the underlying neuronal circuitry with single cell-resolution (see ref^13^ for a recent overview) will be challenging^14^ and remains unsolved. Imaging genetically encoded activity indicators using two- and three-photon based excitation enables a link to be made between neuronal activity and genetically defined neuronal subtypes^15^, reconstructed neural circuitry^16–19^ and when used in miniaturized head-mounted microscopes, to freely moving behavior^20^. Two-photon excitation-based^21^ head mounted microscopes^20,22,23^ enable planar imaging of large neuronal populations^24^, neuronal substructures^23^, and recent developments to miniature microscopes have enabled volumetric and multiplane scanning^25,26^ utilizing electric tunable lenses (ETLs)^27^. Because of the inherent depth limit^28^ of two-photon excitation (2PE) based imaging in densely labelled tissue^29^, this volumetric imaging approach is limited to the upper cortical layers, with the depth-range limited by the ETL axial imaging range and the and temporal resolution limited by the slow duty cycle when sequentially scanning (∼3Hz duty cycle over 180 μm). More recently, the implementation of three-photon excitation (3PE) based fluorescent microscopy^30–32^ to head-mounted microscopes^33–35^ has enabled activity recordings from the deepest cortical layers for extended time-periods from mice freely moving between lit and dark environments^34^. The excitation light properties required for 2PE and 3PE-based imaging of fluorescent activity-indicators, brain-tissue scattering properties, attenuation from water absorption, and tissue damage from heating make 2PE well suited to upper cortical layers, while 3PE allows functional imaging in deeper cortical layers^29^, with an optimal cross-over depth of ∼450 μm from the surface^36^. Here we combined 2PE and 3PE^37^ with temporal multiplexing^38^ to create five imaging planes vertically separated across superficial and deep cortical layers, allowing functional imaging from >1800 neurons per imaging session, for weeks, in freely behaving mice. Because excitation light was delivered through a single fiber for each imaging plane, enabling near-simultaneous (8ns between planes) imaging from the five vertically separated planes^39^, we could compare simultaneously collected activity patterns of layer 2/3 and layer 5 populations from the posterior parietal cortex in mice performing gap-crossing decision making tasks.

## Results

### Head mounted multiplane microscope enables neuronal activity imaging simultaneously across multiple cortical layers in freely moving mice

To create a lightweight head-mounted microscope with multiple, vertically separated imaging planes for freely behaving mice, we developed a system for delivery of excitation light through multiple optical fibers with the fiber tips positioned at different distances from the collimation lens^39^, enabling spatially separated focal points for each fiber after the objective (Fig. 1a, Suppl. Fig. 1). To enable simultaneous imaging from neuronal populations distributed throughout the cortical thickness, we used a laser source providing synchronized pulses at two wavelengths for imaging with both two- (2PE, 960nm) and three-photon excitation (3PE, 1300nm) of GCaMP-based activity indicators^15,40^.

**Figure 1.**
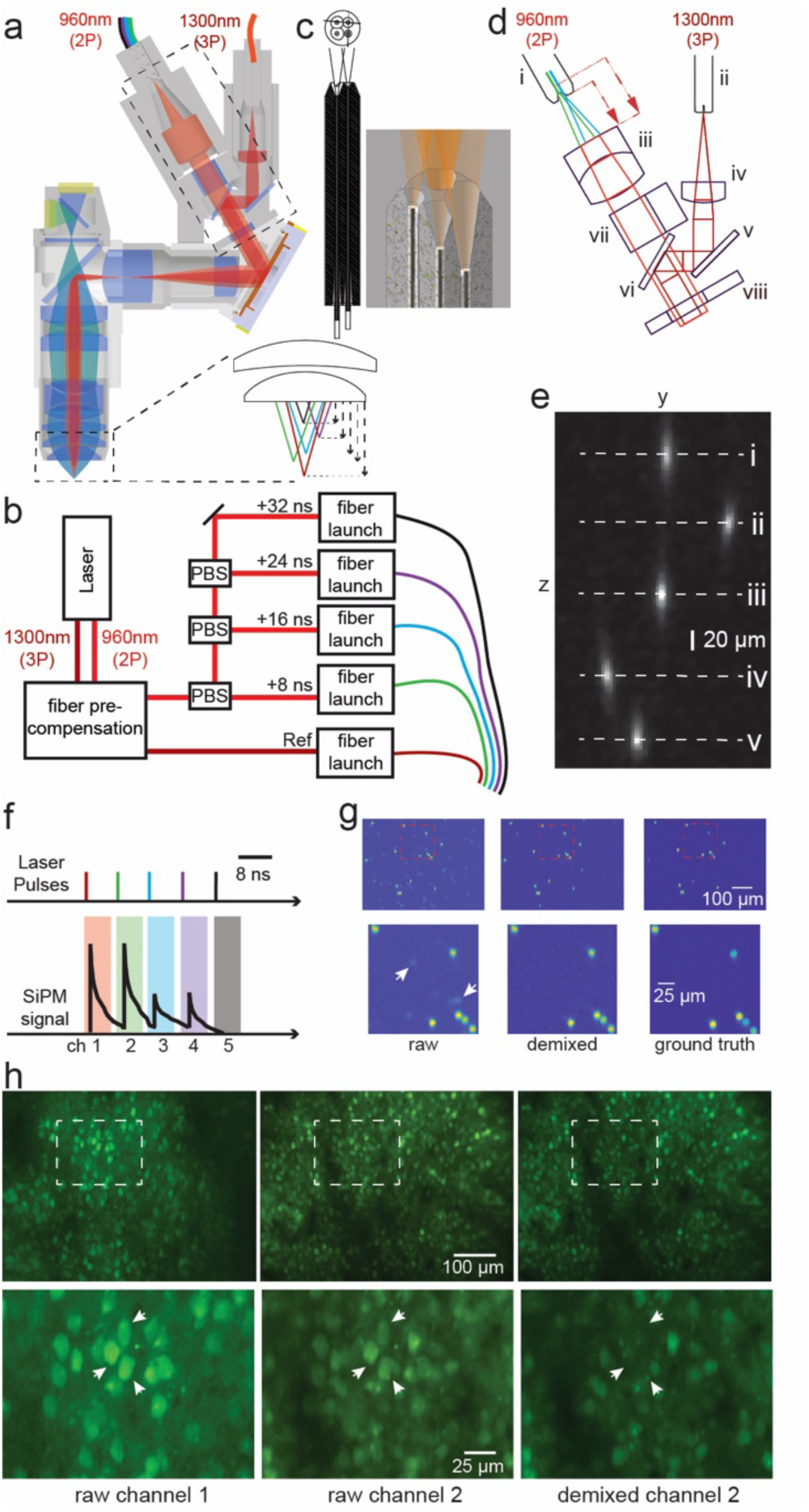
A lightweight head-mounted multiplane fiberscope for imaging across cortical layers in freely moving mice. **a**, microscope schematic showing the 2PE and 3PE fibers, ferrule arrangement and optics used to achieve excitation of 5 imaging planes with axial offset (component list in Suppl. figure 2). **b**, schematic layout of the optical table with the synchronized laser source for 960 nm and 1300 nm, delay structure and coupling to fibers. PBS: polarizing beam splitter. **c**, detail of CAD design of the 3D printed ferrule for the four 2PE fibers. **d**, optical layout of 2PE and 3PE microscope input paths (dashed box in a), showing the 2PE and 3PE ferrules (i & ii), two example 2PE beams (cyan and green), the 3PE beam (dark red), their respective collimating lenses (iii & iv), the mirrors used to combine them into the same beam-path (v & vi) and the ETLs (vii & viii). The distance from each fiber tip to the collimating lens and the lens effective focal length determine the axial offset between imaging planes. **e**, side-view of a 1 mM sulforhodamine solution showing the volumes excited by the beams from all fibers (2PE: i, ii, iii & iv, 3PE: v). Note that the 1300nm 3PE beam excites sulforhodamine by 2-photon excitation. **f**, schematic illustration of the timing of multiplexed pulses (upper), the corresponding gating for signal de-multiplexing (lower, colored boxes) and the SiPM signal (lower black). **g**, average projection of a z-stack through a sample of 10 µm fluorescent beads imaged with laser power on all fibers (top left), the same data after demixing (top middle) and the same sample imaged with laser power on only one fiber (top right). Lower images show enlarged the regions highlighted with red-dashed boxes in the upper panels. White arrows show contamination from other channels visible in the raw, but not de-mixed or ground truth images. **h**, example average images of GCaMP7s-labelled neurons in primary visual cortex of a freely-moving mouse from a leading channel (upper left), a trailing channel (upper middle) and the same trailing channel after de-mixing (upper right). Lower images show the regions outlined with white-dashed boxes in the upper panels enlarged. White arrows show neurons in the image from the leading channel (lower left) contaminating the trailing channel raw data (lower middle), removed after de-mixing (lower right). All images average of 6000 frames. Scale bars under upper and lower images apply to all images in the respective row.

The microscope described here has four 2PE imaging planes and one 3PE plane. The synchronized 2PE and 3PE pulses allowed temporal multiplexing of excitation sources^38^ to enable simultaneous (in the biological time frame) imaging of all planes (8 ns per pixel per imaging plane, 32 ns duty cycle per pixel for all planes). The 2PE beam was split sequentially into four channels using half-wave plates and polarization-dependent beam splitters for adjustable splitting ratio, which allowed the pulse energy to be manually adjusted for each excitation channel. The pulse arrival time for each 2P excitation channel was staggered by adjusting the optical path lengths of each beam to introduce a +8 ns delay between successive excitation channels, using the 3PE channel as reference (Fig. 1b). Dispersion compensation tailored for the 2PE^20,23^ and 3PE^33^ beam was incorporated as the first step in each excitation pathway and prior to splitting and temporally-staggering the 2PE beams. Each beam was then delivered to the microscope using individual fiber launchers and hollow core fibers, with the final excitation consisting of one 3PE and four 2PE time-multiplexed beams (Fig. 1b). To reach optimal excitation efficiency and pre-compensation, each fiber was aligned with respect to the orientation of the polarization of each laser beam (see Methods). To enable precise positioning of the 2PE fibers relative to the microscopes optical path, we designed a single custom-made ferrule with defined 3D positions for each of the four bare fiber-tips (Fig. 1c). The custom ferrule was 3D printed using a high-resolution stereo-lithography printer enabling the 3D relative positions of the fiber tips to determine the relative focusing position of all four 2PE beams. The fiber tip positions were optimized to minimize lateral displacement of the respective focal points (and by extension imaging planes), while introducing sufficient separation between the fiber tips to provide 80 µm axial spacing between two adjacent focal points. The lateral distance between fibers was made as small as possible, and was ultimately limited by the thickness of the protective coating of the fibers. Axial spacing of 80 µm between the 2PE imaging planes was chosen to reduce optical cross-talk between adjacent planes due to the extended axial point spread function (PSF), and provided a total axial span of 240 µm sampled by the four 2PE focal volumes. The 3PE fiber was held in a separate ferrule to allow independent adjustment of the 3PE focal point relative to the four 2PE foci, with the 3PE and 2PE excitation paths being combined using two dichroic mirrors (Fig. 1d, Suppl. Fig. 1b,c, Suppl. Fig. 2). To confirm the axial separation of the focal points from the five excitatory beams, we generated PSFs for all channels in a 1 mM solution of sulforhodamine (101) while imaging the resulting fluorescence with an orthogonally-mounted camera (Fig. 1e). The five spatially-separated PSFs, corresponding to the five excitatory paths present in the microscope, were independent, as verified by sequential blocking of each laser beam (Suppl. Movie 1, microscope resolution further described below). Scanning the 2PE and 3PE focal points in 2D using a MEMS scanner provided five spatially separated imaging planes, with the emitted fluorescence from all planes being split into green and blue color channels using a dichroic mirror and detected by two on-board silicon photomultipliers (SiPM). The time multiplexing introduced into the excitation laser pulses was then used to define temporally separated integration windows for each channel (Fig. 1f), with the high bandwidth of the SiPMs allowing the de-multiplexing of the signal into five separate planes using a single SiPM per color channel (Fig. 1f). To introduce remote focusing functionality, two electrically-tunable lenses (ETLs) were incorporated into the microscope design (Suppl. Fig. 2), one adjusting the position of all imaging planes, and the other allowing independent positioning of only the 2PE planes relative to the 3PE plane. As ETLs in miniature microscopes^24^ are typically not used in a configuration where they are conjugated to the back focal plane of the objective lens, the excitation NA and the optical resolution of the microscope is not constant over the full tuning range. Here they were incorporated for the purpose of fine adjustment of the imaging depth, mainly for the retrieval of same neuronal populations over successive imaging sessions, with functional imaging and characterization of the microscopes resolution all done with the ETLs set close to the center of their range. The maximum field of view of the microscope was 450 x 600 µm², scanned at 9.73 Hz with pixel resolution of 410 x 340 pixels. The lateral and axial optical resolution of the microscope in the configuration described above was measured to be respectively 1.30 ± 0.03 µm and 14.8 ± 1.1 µm for the 3PE plane and 1.45 ± 0.11 µm and 22.2 ± 1.8 µm for the 2PE (mean ± SD, all measured with a sample of 1 µm diameter fluorescent beads, Suppl. Fig. 3).

### Electronic cross-talk removal

The ideal case for de-multiplexing detected fluorescence signals has fast rising and short-lived excitation pulse-evoked photon transients which have fully decayed before the start of the integration window for the next channel. In the current miniature multiplane microscope the detected fluorescence signal was transmitted to a remote high-speed trans-impedance amplifier over 2 m long coaxial cables (see Methods). The combination of the SiPM detector behavior and long transmission lines resulted in slower photon transients than can be achieved with equivalent size-unlimited large photo multiplier tubes^37^, resulting in a pronounced contamination in trailing channels from leading channels (Fig. 1g,h). Multiplexing cross-talk between channels was characterized in the current microscope by imaging 10 µm diameter fluorescent beads (Fig. 1g, Suppl. Fig. 4a). Signals consistent with multiplexing cross-talk were present in the later detection channels (Fig. 1g and Suppl. Fig. 4a-c). As the multiplexing cross-talk was unidirectional, originating from earlier channels only (Suppl. Fig. 4), we developed an iterative optimization routine that minimized this cross-talk between two channels by subtracting a fraction of the intensity of the leading channel from the trailing channel (see Methods). To measure the accuracy of this de-mixing approach we generated ground-truth data by sequentially imaging a volume of fluorescent beads with all channels operating and then imaging the same volume with the same microscope but with only one channel operational (Fig. 1g, Suppl. Fig. 4a). Average fluorescence in regions of interest (ROIs) defined in the leading channels but computed in trailing channels (see Methods) were reduced by a factor of 4.4 ± 1.8 (mean ± SD) after cross-talk removal, and were not significantly different from same measurement on ground-truth data using the same ROIs (Wilcoxon’s rank sum test, Raw vs Ground-Truth p-value 0.0286, Demixed vs Ground-Truth p-value 0.2). Multiplexing cross-talk was also present in *in vivo* images of neuronal populations expressing GCaMP7s (Fig 1h) and using the same de-mixing approach was effectively removed for each channel (Fig. 1h). All following descriptions and quantification of imaging data were performed on data after de-mixing to remove multiplexing cross-talk.

To establish the utility of the head-mounted multiplane microscope for imaging neuronal activity simultaneously from multiple planes, we next imaged neuronal populations in layers 2/3 and 5 expressing GCaMP7s in freely moving mice (Fig. 2a-c, N = 4 sessions from 3 mice spread over 1 to 4 different days). The combination of microscope, head-mounted implant and mounting system design resulted in stable imaging during behavior (median ± SD Euclidean image displacement over all planes, 0.65 ± 2.74 µm, n= 570000 frames total from 4 sessions from 3 mice, Suppl. Movie 2), with 81.8% of the data having frame-wise image displacement less than 2 µm. During these recordings, Ca^2+^ transients were present in labelled somas and dendrites located in all imaging planes (Fig. 2b & c).

**Figure 2.**
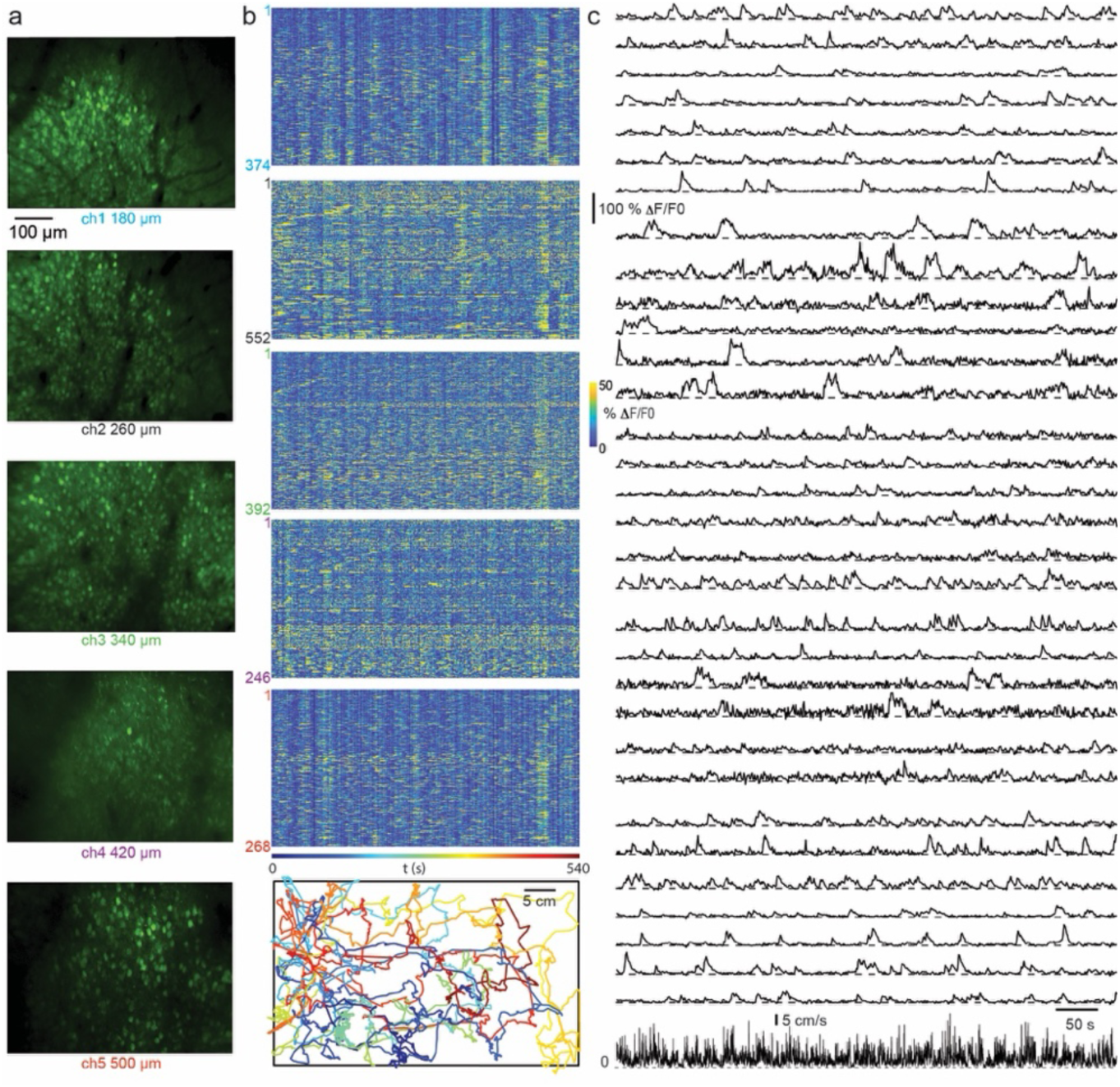
Multiplane imaging of neuronal populations in superficial and deep cortical layers in a freely moving mouse. **a**, average images for each imaging plane of neuronal populations in visual cortex of a freely moving mouse over a ∼10 min long imaging session with associated imaging depths. **b**, raster representations of fluorescence dynamics corresponding to the imaging planes in a. Row show data from identified ROIs. Color scale right of the third panel applies to all rasters. Fluorescence data smoothed with a moving 10 frame average. **c**, animal trajectory in the arena corresponding with the fluorescence data shown in b, color-coded by time. Color scale under lowest raster in b., **d**, example fluorescence traces from the neuronal populations shown in a and b. Bottom trace shows animal velocity.

### Stable imaging from the same neuronal populations across weeks

The total number of neurons imaged by all image planes depended strongly on the homogeneity and density of labelling across the cortical layers. In the experiments described here the number of neurons per imaging plane ranged from 123 to 552, with the average total number of neurons across all imaging planes being 1225 ± 439 neurons per experiment (mean ± SD, range 782 to 1832, n=4 datasets from 3 mice). To assess both the stability and the effects of the multiplane microscope on the tissue being imaged, we first imaged similar neurons (N = 1143) from all imaging planes on two different days (Fig. 3a & b).

**Figure 3.**
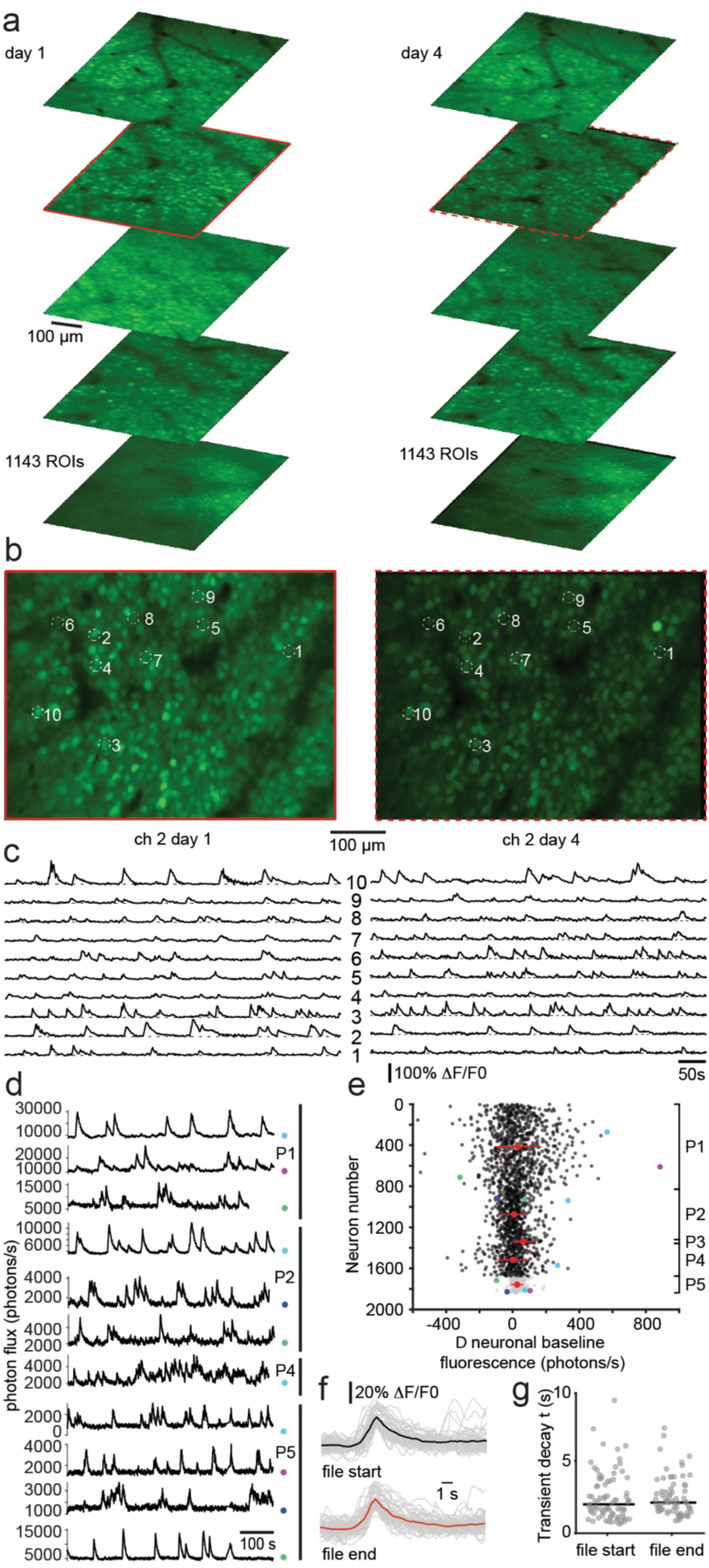
Multiplane imaging is stable over imaging sessions and multiple days. **a**, overview average images from all planes of labelled neurons in the posterior parietal cortex on one day of imaging (left) and near identical populations imaged four days later (right). Both datasets have 1143 identified ROIs. **b**, overview images from channel two in a, showing 10 example ROIs from which fluorescence traces are shown in c. **c** fluorescence traces corresponding to the ROIs labelled in b. **d**, example raw data traces from three 2PE planes (upper, P1, P2 and P4) and the 3PE plane (lower, P5). Color marks denote corresponding points in e for each plane. **e**, scatter plot of the difference in baseline (inactive) raw fluorescence in the first and last 20% of each imaging session for all neurons in all planes (2PE, black, 3PE, gray). Data from individual planes (P1 to P5) indicated on the right. Colored points denote points corresponding to the example traces shown in panel d. Red circles and bars show median and standard deviation for each plane. Data from 4899 neurons, from 4 datasets from 4 mice. **f**, example transients from the first (upper) and last (lower) 20% of recordings used for the analysis of transient decay time constant in e (n=50 transients in both cases, mean trace for first and last examples shown in black and red respectively). h, decay time constants (τ) calculated from exponential fits to individual transients. Data from 81 and 70 individual transients for first and last 20% of recordings respectively. Black bars show median.

Overview images from all imaging planes allowed unambiguous matching of neurons in the two datasets, which were acquired three days apart (Fig. 3b). Similar patterns of Ca^2+^ transients were recorded in the same neurons in both datasets (Fig. 3c). In a second dataset, similar neuronal populations were imaged on two occasions separated by 18 days, with vigorous activity measured from the populations on both days (Suppl. Fig. 5). Together this suggests that the approx. 10- minute-long sessions of continuous imaging had minimal if any detrimental effects on the individual neurons. Both the resting (inactive) neuronal fluorescence and neuronal Ca^2+^ transient decay time constant increase after photodamage caused by excessive exposure to multiphoton excitation sources^41,42^. To quantify the influence of the multiplane microscope on the tissue during free behavior we therefore measured both of these quantities and compared their values at the beginning and end of the imaging sessions. There was no significant difference for any of the imaging planes between resting neuronal fluorescence at the start or end of a recording (Fig. 3d & e, neuronal median ± SD photon flux, calculated over the first and last 20% of the recording respectively, plane 1, 337.0 ± 2003.3 photons/s, 370.8 ± 1984.8 photons/s, p = 0.69, n = 827 neurons; plane 2, 649.6 ± 612.8 photons/s, 638.8 ± 618.6 photons/s, p = 0.92, n = 493 neurons; plane 3, 971.4 ± 692.8 photons/s, 1010.0 ± 708.6 photons/s, p = 0.62, n = 37 neurons; plane 4, 623.2 ± 923.6 photons/s 630.2 ± 951.0 photons/s, p = 0.90, n = 319 neurons; plane 5, 366.1 ± 465.5 photons/s, 394.6 ± 467.3 photons/s, p = 0.30, n = 162 neurons; plane 1 most superficial, N=4 datasets from 3 mice, Wilkoxon’s rank sum test in all cases) nor was there a significant difference in transient decay time constant (median ± SD transient decay τ, start, 2.052 ± 1.718 s; end, 2.188 ± 1.406 s; n=81 and 70 transients respectively, p = 0.36, Wilcoxon’s rank sum test, data from 4 datasets from 3 mice). This indicates that the multiplane microscope does not induce significant photodamage, even during sustained and repeated imaging sessions. Movement of the animals was also not impaired while carrying the microscope, with mice actively exploring the behavioral arena and readily crossing a gap between two elevated platforms. Average velocity during imaging recordings measured for the mice while running was 10.0 ± 6.0 cm/s (average recording time 577.6 ± 53.6 s, n=2 recordings from 2 mice), which is similar to previous measurements made with a previous headmounted microscope^34^ (9.0 ± 0.85 cm/s) and to measurements made from mice carrying only LEDs for position tracking^34^ (10.72 ± 1.75 cm/s).

### Linking decision making free behavior to neuronal activity patterns across cortical layers

We next measured neuronal activity simultaneously from cortical layer 2/3 and 5 in the posterior parietal cortex of mice during complex behaviors (Fig 4a & b, Suppl. Fig. 6). Mice were trained to estimate the distance of a gap between two raised platforms and subsequently cross from one platform to the other to receive a food reward (Fig. 4c).

**Figure 4.**
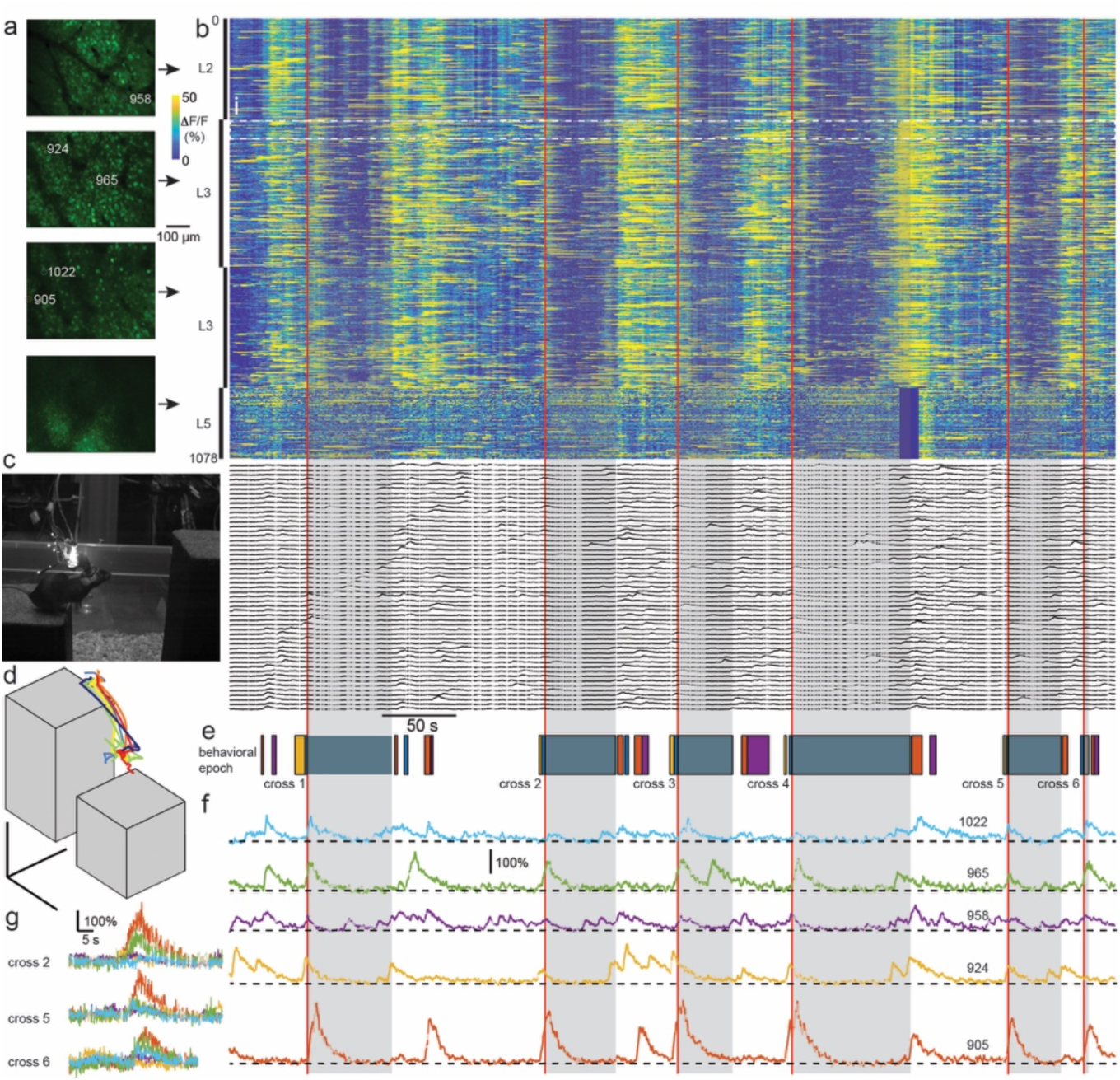
Simultaneous imaging of neuronal activity across layers in posterior parietal cortex during repetitive complex gap crossing behavior. **a**, overview images of neuronal populations in posterior parietal cortex from 4 planes (uppermost in layer 2, second and third from top in layer 3 and bottom in layer 5) during the gap-crossing session in which the data in panels b-g were recorded. **b**, raster representation of 1078 ROIs defined in all 4 imaging planes shown in a. Red lines show cross times. Black arrows indicate correspondence between overview average images in a and the relevant section of the raster representation. Dashed white box shows the 53 neurons for which fluorescence traces are shown below the raster. **c**, image of the mouse just before jumping. **d,** measured 3D trajectories for all identified gap crosses in the imaging session shown in a,b,e & f. Colors for cross 1-6 respectively dark blue, light blue, green, yellow, orange and red. **e**, temporal representation of identified behavioral events in the recording session (blue, cross to top platform, orange, cross to middle platform, magenta, cross from middle platform to arena floor, gray, reward location on top platform). **f**, fluorescence traces of 5 example neurons with transients around gap crosses. **g**, overlay of fluorescent traces from gap crosses 2, 5 and 6 for the same neurons as in f.

Neuronal populations were recorded in ∼10 min. long files enabling calcium transients to be continuously recorded from large populations spread across the cortical layers during multiple gap cross-attempts (Fig. 4d, mean ± SD file duration, 608.2 ± 9.6 s, sessions of 1-2 imaging files, n = 4 datasets from 2 mice). Through a combination of behavior tracking and behavioral monitoring with visible light cameras we could accurately determine the animal’s position in the arena (Fig. 4d) as well as make classifications of the animal’s behavior (Fig. 4e) throughout the task. From these classifications, (see Methods) we established a common behavioral point of reference for each of the trials to correlate neuronal activity to each of the behavior epochs (Fig. 4e & f). As posterior parietal cortex layer 2/3 neurons display repetitive neuronal activity patterns correlated with the various epochs of decision making behavior in mice engaged in virtual reality-based tasks^43^ as well as in freely navigating rats^44^ we next analyzed the neuronal kinetic traces from all planes synchronized to the time that the animal landed on the target landing platform (Fig. 4f). This analysis showed that a subpopulation of layer 2/3 neurons were consistently active before, during, and after the gap-crossing-landing time for each gap-crossing trial (Fig. 4g). As the previous imaging studies have been limited to layer 2/3 of head fixed mice without the access to layer 5 neurons, we next compared the activity patterns of layer 2/3 to those simultaneously recorded in layer 5.

### Differential timing of neuronal activity patterns between the cortical layers during decision making

Ranking all neurons by the timing of the onset of their activity relative to the behavioral point of reference, in this case time of contact to the target platform, for individual gap-crossing trials (Fig. 5a) we established which neuronal subpopulations in each layer were active at a consistent time point during the behavioral sequence (Fig. 5b). Comparing the activity between cortical layers, we found that compared to L2/3 neurons, a smaller fraction of layer 5 neurons were active before and during the crossing events (Fig 5b).

**Figure 5.**
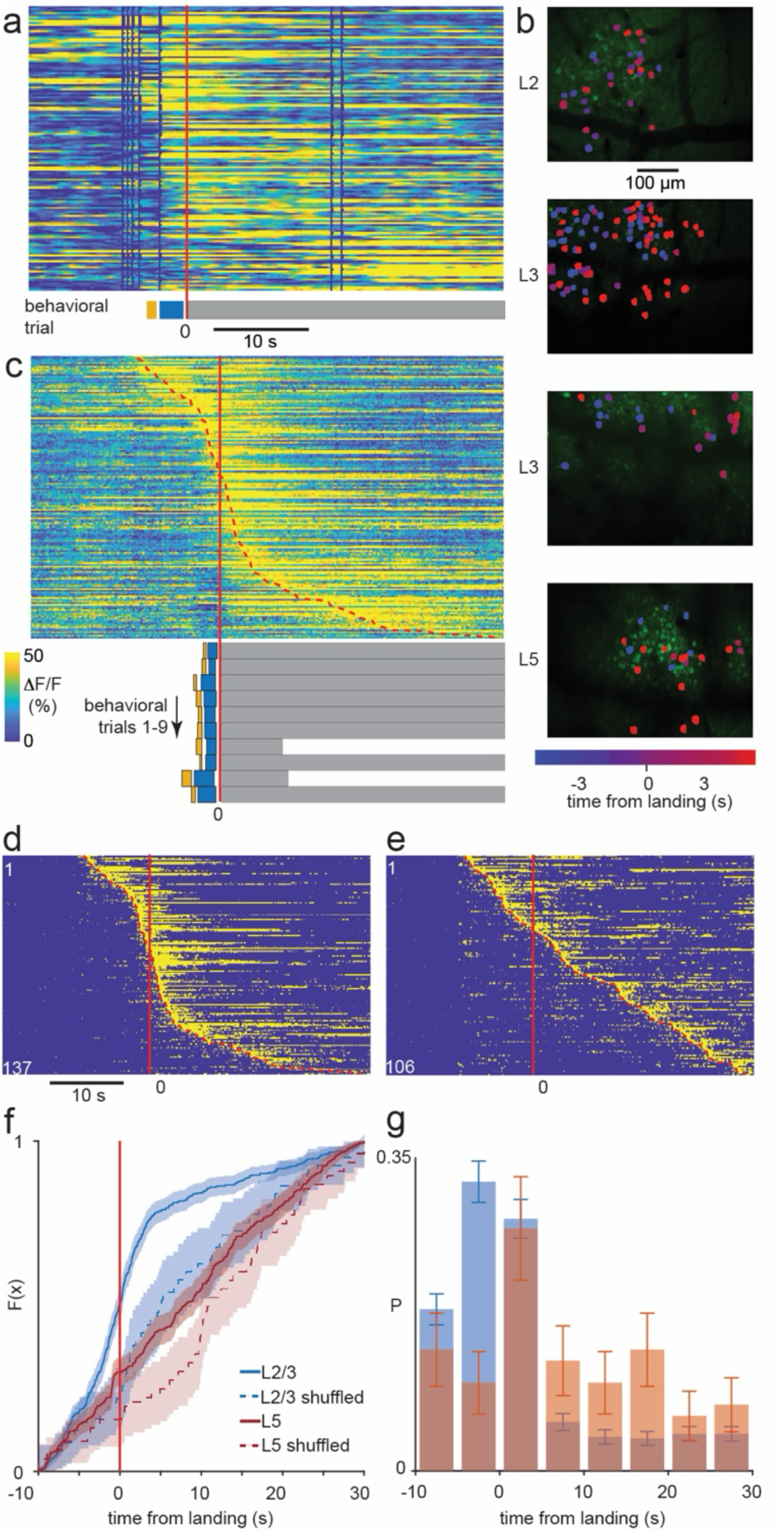
Differential neuronal activity sequence patterns between posterior parietal cortical L2/3 and L5 during decision making. **a**, raster representation, for an example single gap cross from one session from one animal, of neuronal fluorescence traces in the window from 20 s before to 30 s after the cross. Traces are only shown for neurons with significant activity onset, in the above time window, in their fluorescence traces averaged over all gap crosses and synchronized on the time of contact with the top (reward) platform (red line). Neurons are ordered by time of activity onset, from earliest at the top to the latest at the bottom. Colored bars under raster show the behavioral classification during the same time window (blue, cross to top platform, orange, cross to middle platform, gray, reward location on top platform).Color scale under raster in c. **b**, overview images for four imaging planes from the same session in which gap cross shown in a occurred. Colored ROIs represent neurons with activity onset in the time around the cross color coded by onset time. Color scale under bottom image. Putative cortical layers according to imaging depth shown left of the images. **c**, as in a, but showing fluorescence traces averaged over all gap crosses in the session (n = 9 trials). Red doted line represents detected onsets of each neuron. Behavioral classification as in a. **d**, binarized display of data shown in c. Solid and dashed red lines as in c. **e**, as in d with timing of gap crosses randomly shuffled. **f**, cumulative probability functions of onsets of imaged neurons in layer 2/3 (red solid) and 5 (blue solid), with corresponding functions from shuffled data for layer 2/3 (dashed red) and 5 (dashed blue). Confidence interval (95%) shown shaded in corresponding color for both traces. Data from all datasets from 3 imaging sessions from 2 animals. **g**, probability density corresponding to the cumulative probability functions in f for L2/3 (blue) and L5 (red). Error bars show standard deviation, assuming Poisson distribution.

To quantify this, we next binarized the neuronal activity (see methods), fitted the activity onset for each neuron in the ranked activity sequence (Fig. 5d) and compared this to the same data but with the neuronal activity temporally shuffled (Fig. 5e). Neuronal activity across theses behavioral periods for both L2/3 and L 5 populations was significantly different to shuffled data (see Methods) around the landing event (N = 2 mice, 3 imaging sessions), with the density of sequence cells in L2/3 on average 2.07 and 2.22 times larger than in shuffled data for time intervals 5-0 s before the gap crosses and 0-5 s after gap-crosses (Wilcoxon’s rank sum test, p-values <0.01 and <0.0001, see Suppl. Fig. 7 for full statistics) and in L5 on average 2.95 times larger than in shuffled data for time intervals 0-5 s after gap-crosses (Wilcoxon’s rank sum test, p-values 0.0182, see Suppl. Fig. 7 for full statistics). In addition. comparing the L2/3 and the L5 neuronal responses during these behavioral epochs showed a significant increase in number of neurons active in sequences before the crossing event in L2/3 but not in L5 (Fig 5f & g, Suppl. Fig. 7). Together this suggests that neuronal activity in the posterior parietal cortex is differential across layers during free behavior, and while the layer 2/3 neuronal activity shows a strong repetitive temporal sequence during the entire behavioral sequence, layer 5 neurons are more active in the post-decision period during reward acquisition.

## Discussion

We designed and implemented a multiplane microscope enabling the measurement of neuronal activity from large cortical populations, spread across the cortical layers, in mice performing complex freely moving behavioral tasks. The microscope was designed to measure from the same neuronal populations over days and to simultaneously measure activity from neuronal populations located in deep and superficial cortical layers. Because there was an 8 ns delay between each plane when delivering the excitation pulse, and one excitation pulse per pixel, the total frame rate was not affected by the number of planes but instead was determined by the MEMS scanner performance, enhancing the global data acquisition limit by a factor equal to the number of fibers. This temporal multiplexed multi-fiber approach provides the advantages, compared with previous multiplane approaches using ETL^24–26^, of enabling the combination of both 3- and 2-photon excitation modalities and their imaging advantages^29,36^ as well as enabling near-simultaneous sampling across the vertically separated planes. In addition, as there were separate ETL’s in the 2- photon and 3-photon excitation pathway, the axial position of the imaging planes for each modality could be independently adjusted, without diminishing the optical properties of the PSF, allowing neuronal populations to be located from the previous imaging sessions. Finally, by combining multiple fibers, with axial offsets, ETLs and 2PE and 3PE, the axial range between imaging planes was stretched to cover the distance between L2 and L5 of mouse cortex, enabling simultaneous imaging of deep and shallow cortical layers.

The high frequency electronics required for the detection system, combined with the characteristics of the connections between the microscope mounted detectors and signal digitization and amplification made it vulnerable to generating ‘ghosting’ artefacts from one imaging channel to the next^37^. We established, using ground truth data, that the magnitude of the contamination in the effected channels was small in comparison to the magnitude of the actual fluorescent structures in the imaging plane^37^. Additionally, as the artefact was temporally unidirectional, we could build a simple removal approach that allowed the effective de-mixing of the signals from the affected channels without any detectable degradation of the imaging.

As the extra excitation delivery fibers added negligible burden and weight combined with our previously designed on-board detector-system^34^ that removed the plastic optical fiber burden and enabled imaging in lit conditions, this current design was well suited to studying complex free behavior, such as gap crossing shown here. While the weight of the new components slightly increased the microscope weight compared to our previous version, in combination with appropriate counter-weighting the weight experienced by the mouse is reduced to the point where they were able to jump across a gap between raised platforms and to move with little to no hindrance during extended imaging sessions (longest here approx. 30 min). In addition, the measured characteristics of animal behavior not significantly different to previous microscope^34^ or when carrying tracking struts alone. One major concern was increasing the amount of energy into the tissue^37^, while the total number of pulses delivered into the tissue was in the same range as our previous microscopes, the combination of both 2- and 3-photon excitation was could potentially create tissue heating and damage^45^. Using commonly described measurements of tissue health^42^ we could not detect any damage associated with excessive energy absorption throughout the 5 imaging planes, even after continuous imaging sessions of >30 mins.

Together, this enabled imaging activity from neuronal located in both layer 5 and layer 2/3 during complex decision-making behaviors where the animal had to estimate the gap distance and cross from one platform to another platform to receive a reward. We chose to image neuronal populations located throughout the posterior parietal cortex during decision-making behaviors in mice, as the correlation between behavior and upper cortical layer neuronal activity is well established^43,46–48^. With the ability to record over many imaging sessions where the animals were free to perform the crossing task at will, we were able to establish the differences between layer 2/3 neuronal activity patterns and layer 5 activity patterns relative to the animal’s behavioral outcome. As this new microscope enables near-simultaneous recording of neuronal activity across cortical layers in animals performing decision making tasks, together with high-resolution circuit reconstruction techniques^16^ questions about how individual neurons located across the cortical layers transform sensory-derived inputs^17^ during free behavior can start to addressed at the circuit level.

## Methods

### Animals and surgical procedures

#### Animal protocol

All animal experiments were conducted in accordance with the animal welfare guidelines of the Max Planck Society and with animal experimentation approval granted by the Landesamt für Natur, Umwelt und Verbraucherschutz Nordrhein-Westfalen, Germany (protocol number 84- 02.04.2020.A403).

Experimental animals were 4 male wild type C57Bl/6 mice (*Mus musculus*) obtained from Charles River, Germany. The mice were housed in a specific-pathogen-free temperature- (21±1°) and humidity-controlled (>45%) facility on a 12 h light–dark cycle with food and water available ad libitum. Mice were group-housed until surgery and singly housed afterwards. At the start of the experiment the mice were between 26 and 31 g (mean 28 g).

For virus injections we used AAV1/2-syn-jGCaMP7f based on the plasmid pGP-AAV-syn-jGCaMP7f- WPRE (plasmid #104488) published in^49^.

#### Surgical procedures for fluorescent labelling of neurons with jGCaMP7f

Prior to surgery, all instruments, including the glass injection capillary, were sterilized by either autoclaving or heat sterilization. Animals were anesthetized with an intraperitoneal injection of a three component anesthetic cocktail (FMX) consisting of fentanyl (50 µg/kg, Hameln pharma plus GmbH, Hameln, Germany), midazolam (5 mg/kg, Hameln pharma plus GmbH, Hameln, Germany) and xylazine (10 mg/kg, WDT, Garbsen, Germany). Body temperature was maintained at 37–37.5 °C with a heating pad and heater controller (FHC, ME, USA). Animal status and depth of anesthesia was monitored approx. every 15–30 min, and anesthesia maintained throughout with supplementary doses of 30–80% of the above anesthetic combination given as necessary in order to maintain absence of withdrawal and corneal reflexes. The animals were then placed in a stereotaxic apparatus, the hair on the scalp removed and the skin cleaned with 70% ethanol. The right parietal bone was exposed and a burrhole drilled (approx. 500 µm diam.) at 3.6 mm posterior and 2.6 mm lateral relative to bregma for experiments in visual cortex (V1) and 2.1 mm posterior and 1.4 mm lateral relative to bregma for experiments in posterior parietal cortex (PPC). A small slit was made in the dura underlying the burrhole, and a glass injection capillary with a beveled tip containing a high titer solution of AAV1/2 coding for jGCaMP7f was advanced into the cortex. For V1 experiments, the pipette was angled at 25° relative to horizontal and was advanced posteriorly along a trajectory parallel with the saggital suture. Three virus injections were made targeted to depths of 600, 500 and 300 µm (tangential to cortical surface) and of volume 80, 50 and 50 µL respectively. The deepest of the three injections with the above burrhole and pipette orientation and advance was targeted to be 4.5 mm posterior and 2.6 mm lateral relative to bregma. For PPC experiments, the pipette was angled at 28-31° relative to horizontal and was advanced posteriorly along a trajectory parallel with the saggital suture. Three virus injections were made targeted to depths of approx. 450, 320 and 200 µm (tangential to cortical surface) each 200 µL in volume. In all cases there was a 5 minute delay between the end of the virus injection and moving the pipette to either the next injection location or withdrawing the pipette from the cortex. After withdrawing the pipette, the craniotomy covered with medical silicone (KwikSil, WPI, FL, USA) and the skin sutured closed using 5/0 vicryl sutures (Ethicon, NJ, USA). Approximately 30 minutes prior to the completion of the procedure the animals were administered buprenorphine (30 µg/kg, Bayer, Leverkusen, Germany) and carprofen (5 mg/kg, Zoetis, NJ, USA) for post-operative analgesia, and after the completion of the procedure a cocktail of antagonists to the anesthetic drugs (anti-3K) consisting of naloxone (11.2 mg/kg, Ratiopharm, Ulm, Germany), flumazenil (0.5 mg/kg, Hikma, Amman Jordan) and atipamezole (10 mg/kg, Orion Pharma, Hamburg, Germany).

#### Surgical procedures and imaging in freely moving animals

All surgical instruments and solutions used were autoclaved prior to commencement of the procedures described below. Three to five weeks after the surgery to label neurons with jGCaMP7f, animals were anaesthetized with the FMX cocktail described above, and body temperature maintained at 37–37.5°C. Animal status and depth of anaesthesia monitoring procedures were as described in the section on labelling neurons with jGCaMP7f. Anesthesia was maintained with supplementary doses of 30–80% of the FMX solution. The hair on the dorsal aspect of the skull was removed and the skin cleaned with 70% ethanol. A midline incision in the skin over the parietal bones was made, the skin retracted and galea removed to expose the parietal bones, including the site of the previous burrhole. The exposed bone was then cleaned with hydrogen peroxide solution (3% by volume in sterile saline) and thoroughly washed with sterile saline. The bone was then mechanically roughened prior to application of a layer of dental adhesive (Optibond, Kerr, CA, USA). A custom-made headplate, with a central circular aperture measuring 3.3 mm in diameter, was then fixed to the skull over the Optibond layer with dental composite (Charisma, Kulzer GmbH, Hanau, Germany). The central aperture was placed such that the burrhole from the previous surgery was located approximately centrally in the medial-lateral axis and near the anterior edge of the aperture. The skin incision was then closed firmly around the headplate using 5/0 vicryl sutures (Ethicon, NJ, USA). A circular craniotomy with a diameter of approx. 3 mm was then opened in the center of the headplate aperture, including at the anterior margin of the site of the previous craniotomy. The dura was removed and the cranial window closed using a pre-formed plug and coverslip (circular, 5 mm diam., 100 µm thickness, CS-5R-0, Warner Instruments Holliston, MA, USA; plug 300 µm tall glass cylinder fixed to the coverslip using UV curing optical glue (Norland Optical adhesive No. 68, Edmund Optics, Mainz Germany)), which was secured to the base of the headplate using transparent biocompatible silicone (KwikSil, WPI, FL, USA). The plug was designed to approximately match the gap between the surface on which the coverslip rested and the bottom of the bone, including the thickness of the headplate and the adhesives used to fix it to the skull. The cranial window was protected using a plastic 3D-printed cap which was secured in place on the headplate using fastening screw. Post-operative analgesia and anesthetic antagonist solution composition, dose and administration were the same as described above in the procedure for labelling of neurons with jGCaMP7f.

The head-plate consisted of a printed component, forming the walls of the head-plate chamber and the stalk for holding it, which was printed in a biocompatible, glass-filled resin (Temporary CB and Form 3 printer, Formlabs, MA, USA), and a 80 µm thick steel ring (6.9 mm diameter with a central aperture measuring 3.3 mm in diameter). The two head-plate components were then fixed together using dental composite (Charisma, Kulzer GmbH, Hanau, Germany).

#### Training gap crossing task

To introduce the animals to the nature of the task and reward, animals were initially placed in a home ‘jumping’ cage (dimensions 55×35.5×30 cm, LxWxH) containing both launch and reward platforms (dimensions below in “*Freely moving experiments*“) with water and/or sweetened cereal pieces as rewards on top of the reward platform. Launch and reward platforms were initially positioned adjacent to one another so that the animals could climb freely between them.

Rewards on the reward platform were replenished regularly to promote frequent exploration and cycling. Once mice were comfortably cycling between blocks (roughly 1-2 days training) the distance separating the launch and reward blocks was increased incrementally until the animals were required to jump in order to obtain the reward. These distances continued to be increased, up to a maximum of 20 cm, or until an individual mouse began refusing to cross more often than it crossed successfully. During periods of water restriction, the daily water allowance was delivered either on the reward platform or during discrete 10–30 minute training sessions, in which each successful cross was accompanied with a water reward of approx. 0.1ml and/or sweetened cereal piece reward. The daily water allowance was determined separately for each mouse, being 50% of ad libertum water consumption by design and a minimum of 30% if required. Finally, to habituate the mice to both the experimental arena and the experimenter, each animal underwent at least three 30-minute acclimation sessions in the laboratory with the microscope and all supporting equipment, using the same platform configuration and arena used in the recording sessions. The mice were allowed to cross freely with each cross rewarded.

#### Placement of the multiplane microscope

Two to four weeks after the procedure to install the cranial window the animals were anesthetized with isoflurane (1.25–2.5%, Baxter, Unterschleißheim, Germany) delivered in air at a flow rate of 1.5 L/min, and transferred to the miniature microscope, which was mounted on a navigation stage for locating an appropriate position for imaging within the cranial window. The navigation stage consisted of a micromanipulator (MP-285, Sutter Instruments, CA, USA) to which the microscope was mounted using a custom-made mount. The mount included two angular kinetic mounts (GN05/M and GN1/M, Thor Labs, NJ, USA) allowing adjustment of tilt with respect to the coverslip and cortical surface. Once a target population of neurons had been located, the intermediate attachment plate (already mounted to the miniature microscope) was attached to the headplate with dental composite (Charisma Flow, Kulzer GmbH, Hanau, Germany). The microscope was then removed, a plastic 3D-printed protective cap attached using the same mounting mechanism used to secure the microscope in place and the animal allowed to recover.

#### Freely moving experiments

Freely moving experiments were subsequently conducted from 9–11 weeks after opening the cranial window. At the commencement of each recording session, the animal was taken from its home cage, the head gently restrained by holding the handle on the back of the headplate, and the microscope placed onto the intermediate attachment plate and secured in place with a holding screw. After microscope placement, the animal was placed on an open rectangular arena measuring 30×50 cm^2^ for experiments in V1, or in PPC experiments into a walled rectangular arena measuring 32×50 cm^2^ with 15 cm tall walls, into which was placed two movable platforms, the reward platform, measuring 10 × 10 × 20.5 cm^3^, and the launch platform, measuring 10×10×12.5 cm^3^ with an attached short step measuring 10×5×6.5 cm (all measurements L×W×H). The mice in the PPC experiments were water restricted as described above. Mice not consuming 30% of their ad libitum consumption in the behavioral arena were given supplementary water making up the difference between their consumption in the arena and the 30% threshold at the end of the session in their home cage. They were further encouraged to cross between platforms using small pieces of breakfast cereal (Froot Loops, Kellog’s, Germany). At the end of a recording session, the animal was gently removed from the arena, the screw on the microscope mounting plate unfastened and the microscope removed, the protective cap put in place and the animal returned to its home cage.

##### Histology

At the termination of the imaging experiments animals were deeply anesthetized with ketamine (100 µg/kg) and medetomidine (200 µg/kg), and perfused transcardially with 0.1 M phosphate buffer (PB) followed by 4% formaldehyde solution (Roti-Histofix, Carl Roth, Karlsruhe, Germany). The brain was then removed, post-fixed at 4°C overnight in the same formaldehyde solution and then transferred to 0.1 M PB. Sections of 100 µm thickness were then cut on a vibrating microtome (Leica VT1000S, Wetzlar, Germany) and mounted in fluoroshield (Sigma, MO, USA). Images were acquired on an inverted microscope (Nikon Ts2R-FL, Tokyo, Japan).

### Hardware

#### Miniature microscope optical configuration and production of the prototype

The optical system of the multiplane microscope, from MEMS scanner to objective and detection system, is largely similar to the previously described miniature multiphoton microscope ^34^, composed of the following components. The scanner is a 2.0 mm diameter MEMS scanner (A7M20.1-2000AU-LCC20-C2TP, Mirrorcle, CA, USA), featuring ±5 mechanical degrees of scanning range and 1.7 kHz scanning frequency. All lenses and mirrors off-the-shelf used were custom- adjusted by centering and thinning down (Throl GmbH, Wetzlar, Germany) to fit mechanical specifications of the microscope (Suppl. Fig. 1). The scan and tube lens consists of a custom doublet lens and a plano-convex (PCX) lens with an effective focal length (EFL) of 12 mm (ø3 mm, (#67-444, NIR II, Edmund Optics, NJ, USA). A dichroic mirror (T875spxrxt, AHF Analysentechnik AG, Tübingen, Germany) was used to separate the excitation beam path and the emitted fluorescence. A condenser lens with an EFL of 5 mm was used (Throl GmbH, Wetzlar, Germany) followed by an infrared filter (G380227032, Qioptiq Photonics GmbH, Goettingen, Germany). A dichroic long-pass mirror (490nm, DMLP490R, Thorlabs, NJ, USA) separated emitted light into two channels, one detecting light emitted from third harmonic generation (THG, blue) and one for GCaMP7f (green).

We used SiPM multi-pixel photon counters (MPPC, S13360-1375PE, Hamamatsu, Hamamatsu City, Japan) as on-board detectors mounted on the microscope.

The optical system between the end of the optical fiber and the MEMS scanner for the 3PE path consisted of a 2 mm diameter, 5 mm EFL aspheric collimation lens (354430-C, Thorlabs, NJ, USA) and two dichroic mirrors (DMSP1180R, Thorlabs, NJ, USA). For the 2PE path, it consisted of a 2 mm diameter, 4 mm EFL achromatic lens (#84-128, NIR II, Edmund Optics, NJ, USA) and a stack of µT lenses (Packaged TLens® Silver IRSM – PIF.15.P01, PoLight ASA, Skoppum, Norway) for adjustment of the axial offset between the 2PE image planes and 3PE imaging plane after the objective lens. An additional single µT lens common for both 2PE and 3PE was inserted before the MEMS scanner for fine axial position adjustment. The full optical system of the miniature microscope is shown in Fig. 1a and Suppl. Fig. 1 and 2, and its various parameters were optimized using OpticStudio 14.2 (Zemax Europe, Bedford, UK).

The production of the prototype has been described in detail in previous work (Klioutchnikov 2022). Briefly, the structure of the microscope body, implants and lens-mounts, including objective, were designed with Inventor (Autodesk GmbH, München, Germany). The parts for the microscope body were printed in black resin V4 (Formlabs, MA, USA) on a 3D printer (Form3, Formlabs, MA, USA). All the optics and the objective were supplied by Throl GmbH (Throl GmbH, Wetzlar, Germany). The four fibers for the 2PE path (HC-920-PM, NKT Photonics A/S, Birkerød, Denmark) were jointly mounted in a 3D printed ferrule (Fig. 1c,d) with a diameter of 1.25 mm, which was printed in yellow resin (HTL -Yellow Resin, BMF Precision, Inc., MA, USA) on a high-resolution stereo-lithography printer (microArch S230, BMF Precision, Inc., MA, USA). The ferrule was then mounted in a corresponding structure in the microscope (Fig. 1a,c&d). The fibers were cleaved and the coating removed with a razor blade over 10–15 mm, then manually inserted in the ferrule and glued inside with UV-curing glue (NOA068, Thorlabs, NJ, USA). A manual fine-tuning of each fiber’s axial position was necessary to achieve the 80 µm offset between the corresponding imaging planes prior to the final gluing of the fibers. For the 3PE path, the fiber was mounted in a similar 3D printed ferrule but designed with a single fiber bore (Fig 1d, Suppl. Fig. 2), with the ferrule then mounted into its corresponding structure on the microscope (Fig. 1a & d). Fiber cleaving and mounting in the ferrule were as described above for the 2PE path.

#### Synchronous multimodality excitation laser setup

For excitation, we used an optical parametric chirped pulse amplifier (OPCPA), pumped by a Ytterbium fiber laser (white dwarf dual, Class 5 photonics GmbH, Hamburg, Germany) that produced 1300 nm pulses with a maximum average power of 4 W (2 µJ at 2 MHz; 3P excitation beam) as well as 960 nm pulses with a maximum average power of 1 W (0.5 µJ at 2 MHz: 2P excitation beam). We used a half-wave plate (AHWP05M-1600, Thorlabs, NJ, USA) mounted on a rotation mount with a resonant motor (ELL14, Thorlabs, NJ, USA) with a polarization beam splitter (PBS104, Thorlabs, NJ, USA) to control beam intensity for the 3P beam and a Pockels’ cell (350-80LA, POLYTEC GmbH, Berlin, Germany) for the 2P beam. In the 3PE path we compensated dispersion introduced by the fiber using the two prism sequence with additional bulk silicon described previously ^33^. In brief, to compensate third-order dispersion (TOD), a double-path two prism sequence compressor with Brewster-angle silicon prisms at 32° apex angle (4155T724, Korth Kristalle GmbH, Altenholz, Germany) was used. The custom-designed second prism had a larger base of 56 mm to increase the compensation range. Additional anomalous group-velocity dispersion (GVD) generated by the prism sequence was compensated using bulk silicon. For pre-compensating the dispersion of the 1.5 m of the hollow-core fiber ^33^ used in this study we used an inter-prism distance of 35 cm with additional 10 cm of bulk silicon. For the 2P excitation, the GVD was minimized by adding 5 mm ZnSe slabs at Brewster angle (WG71050, Thorlabs, NJ, USA), altogether a total path of 16 cm, to compensate for ∼1.75 m of fiber (HC-920-PM, NKT Photonics A/S, Birkerød, Denmark). The beams were coupled into their respective fibers using a range of achromatic coupling lenses with EFLs from 7.5 mm to 20 mm (AC050-XXX-C-ML, Thorlabs, NJ, USA), with the choice individually made by the best coupling efficiency.

To achieve the timing structure of the laser pulses allowing for multiplexed excitation of fluorophores we used cascaded pairs of half-waveplates (AHWP10M-980, Thorlabs, NJ, USA) and polarizing beam splitters (PBS105, Thorlabs, NJ, USA), each pair splitting laser pulses in two and allowing continuous control of the power splitting ratio. After each splitting step, the reflected beam was then coupled into a fiber and a ∼ 8 ns delay was added to the transmitted beam to achieve the required time separation between successive pulses fitting the designed demultiplexing strategy. Each fiber symmetry axis was aligned to the axis of the linear polarization of each incident beam for optimal transmission and temporal profile. For the last fiber the transmitted beam of the third beam-splitter was used. Each pulse propagated then down its respective fiber and excited a volume of fluorophore at a respective plane (Fig. 1a-d, Suppl. Fig. 2). The splitting ratio was manually adjusted for each animal to achieve similar level of excitation for each channel. All four 2PE channels were then jointly modulated with the common Pockels cell for power adjustment, flyback and fill fraction blancking. The achieved time-structure of the multiplexed excitation beam was verified using a fast photo-diode (DET10A2, Thorlabs, NJ, USA).

#### Fluorescence detection, control electronics and software

Scanimage software and high-speed analog to digital and digital to analog converter board (ScanImage®2023.1.0 premium, hsViDAQ, MBF Bioscience, VT, USA) was used to control the microscope and acquire the data. The MEMS scanner was operated in the resonance mode in the fast axis at 1.7 kHz providing 3400 lines per second using bi-directional scanning and 3 ms flyback. This, combined with a 2 MHz excitation laser pulse rate and 80% of imaging fill fraction, resulted in a resolution of about 410 pixels × 340 lines at 9.73 Hz frame rate. The field of view during data collection measured 600 × 450 µm². The driving signals from Scanimage software were sent to the MEMS driver (BDQ_PicoAmp_4.6, Mirrorcle, CA, USA). A high bandwidth preamplifier was custom designed based on the publicly available schematics from the evaluation board of the SiPM (S13360- 1375PE, Hamamatsu, Hamamatsu City, Japan). Gating of acquisition to separate the sequential pulse sequence was achieved by adjusting the FPGA parameters using ScanImage 5 built-in features.

#### Environment light

The environment of the track was homogeneously illuminated using six 24 V RGBW LED strips of 125 cm length with 910 lm/m and eight 12 V white LED strips of 125 cm length with 700 lm/m (both LED strips from PowerLED, Berkshire, United Kingdom), arranged equidistantly in a patch of 125x60 cm^2^ at a distance of 150 cm above the track.

The strips were switched on and off in synchrony with the line signal of the microscope using a custom circuit based on TIP120 Darlington transistors.

#### *Tracking* animal position

Animal position in the arena was tracked using a system of overhead cameras and position tracking LEDs mounted onto the microscope previously described in ^50^. For mounting the LEDs onto the microscope, three struts were mounted onto the microscope body, with three infrared (IR) LEDs (940 nm, SFH 4053, Osram) mounted uniquely per strut. Images of the IR-LEDs were acquired at 200 Hz by four calibrated and synchronized digital cameras (acA1300-200um, Basler AG, Ahrensburg, Germany) mounted above the arena. The cameras were equipped with 4.4–11 mm, F1.6 objective lenses (LMVZ4411 1/1.8” F1.6/4.4–11mm, Kowa, Japan) to facilitate coverage of the arena and IR bandpass filters (BN 940-43, Midwest optical systems, IL, USA) to facilitate tracking of the LEDs only. The exposure active signals from the overhead tracking cameras as well as the frame synchronization signal from the miniature microscope were fed into an analog-digital converter (Power 1401, Cambridge electronic design, Cambridge, UK) and recorded with Spike2 software for synchronization of behavioral tracking and multiphoton imaging data. Animal position and head orientation were tracked as described in^50^, with the exception of two datasets where the positions of the IR LEDs in the camera images were tracked using DeepLabCut^51,52^ and the detected positions then used for determination of animal pose and position as described in ^50^. All cameras were calibrated using the calibration procedure described in ^53^.

### Data analysis

Raw imaging data were first pre-processed to minimize scan-pattern distortions, brain motion and cross-talk between gated channels as described below. The photon-count related signal structure of the SiPM’s used on the microscope allowed all data pre-processing steps to employ photon-flux as the measurement unit, which was determined in the initial photon calibration step. Expected emission photon counts per pixel are on the order of 1 to 10^54^.

#### Photon calibration

The pulse-height distribution of the SiPM’s used exhibited a prominent peak structure related to n- photon detection events. To infer the scaling factor from ADC signal level to photon-count we first characterized the Gaussian peak and width parameters of the noise floor. We then identified the 1- to n-photon peaks using a moving window correlation with a Gaussian with parameters of the noise floor. The pulse-height to n-photon scaling factor was then defined as the average of the difference between the 1- to n-photon peaks and the 0-photon offset was inferred by subtracting the scaling factor from the 1-photon offset. These parameters were then applied to the image sequence using floating-point arithmetic.

#### Cross-talk demixing

The primary sources of cross-talk that we identified were firstly the time course of fluorescent decay and the analog high-frequency behavior of the SiPM detector, cables and pre-amplifier, both having an impact on the scale of ∼10 ns, and secondly after-pulsing of the SiPMs (spurious pulses elicited by the SiPMs after a photon or dark capture event identical in amplitude and shape to the original pulses), which occurs with a delay of 10 to several 100 ns. As the excitation delay between channels is 8 ns, both sources can result in spurious pixel intensities in trailing channels (time range ∼32 ns), but not in next pixel (time range 500 ns) or next frame (time range 100 ms) The net result is the appearance in the trailing channels image of dim replicas of labelled structures from the leading channel at the exact same locations as in the leading channel.

To remove the cross-talk between leading and trailing channels, we determined a cross-talk coefficient (see below) for each pair of leading and trailing channels which could be applied uniformly between the channel pair. Specifically, we applied the linear cross-talk matrix

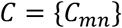

with leading and trailing channels denoted as 𝑛 and 𝑚, respectively, which only contained non-zero coefficients for 𝑚 > 𝑛, so that each channel-wise image series 𝐼_"_ (each image series having identical width, height and frame count) could be de-mixed using

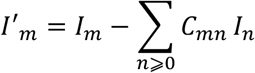

To determine the cross-talk coefficient for each relevant channel pair we first calculated the selected-pixel mean correlation coefficient (spMCC) to estimate the extent of the cross-talk between leading and trailing channel. To reduce the influence of regions in the image lacking labelled structures we identified all non-saturated pixels (pixels with a maximum photon flux < 30) for which the standard deviation of the low-pass-filtered data (Gaussian low-pass, σ ≈ 2.5) was greater than 0.5, then calculated the spMCC as the average of the pixel-wise Pearson’s correlation coefficients between leading and trailing channels for the selected pixels. The cross-talk coefficient for each relevant channel pair was then determined by minimizing the spMCC.

#### Motion correction and scan-pattern correction

For optimal motion correction performance the physical pixel size must be constant over the image, which with a sinusoidal drive signal to the MEMS scanner is not the case. To correct this, we used non-linear resampling over the fast axis. To avoid aliasing during that process, we first upsampled the fast axis tenfold then non-linearly downsample it for spatial uniformity using the resample Matlab function with default parameters for low-pass filtering. For convenience we chose the final pixel resolution to be 512*512 for further analysis, as a balance between more pixels for higher resolution ROI definition and precision of motion-correction, but limited increase in data size and processing time. Motion correction was performed using a custom Matlab port of the image registration component of suite2p^55^, involving a displacement estimation using phase-correlation and subsequent rigid frame shift. To increase robustness under low photon-count conditions (where dark counts interfered with the phase-correlation) the image series was filtered in time (Gaussian low-pass, σ ≈ 1.5).

#### Extraction of fluorescence signal

The fluorescence signal was calculated for regions of interest (ROIs) by averaging pixel values per frame and ROI. Alternatively, to yield photon flux pixel values were summed up. ROIs were manually selected, using custom software, by drawing a line pattern around beads, neuronal somata or dendrites.

#### Calculation of ΔF/F0

The baseline fluorescence 𝐹_$_ was defined as the mean of the lowest 20% of fluorescence values within a window of maximally 200 s around the current frame (if not truncated by beginning or end of the file) after applying a Gaussian filter with a standard deviation of 282 ms.

#### Point Spread Function (PSF)

The PSF was measured using a sample of 500 nm fluorescent beads (F8813, Molecular Probes Inc, OR, USA) suspended in an agarose gel, using image stacks with 1 µm z-step. Average measured optical resolution axial 22.2 ± 1.8 µm, lateral 1.45 ± 0.11 µm for 2PE channels and axial 14.8 ± 1.1 µm, lateral 1.30 ± 0.03 µm (all mean ± SD).

#### Side-view imaging of fluorescent agarose gel

To observe the relative position of the laser spots corresponding to each excitation beam, we placed the microscope above a fluorescent agarose gel filled glass cuvette. A side-view camera with a high-resolution objective lens (MVL25TM23, Thorlabs, NJ, USA) was used to record the fluorescence resulting from the excitation of the fluorophore from the side, see Suppl. Movie 1.

#### Ground truth data for de-mixing algorithm

To provide ground truth data for the multiplexed channels de-mixing algorithm we have acquired sequential images in axial dimension of 10 µm fluorescent beads (F8836, Molecular Probes Inc, OR, USA) suspended in an agarose gel. The imaging was first performed with all excitation beams operating, then the same volume was imaged sequentially using only one fiber at each acquisition. For the latter acquisition all unused laser beams were physically blocked.

#### Quantification of de-mixing algorithm performance

To quantify the performance of the de-mixing algorithm first rescaled the ground truth data described above from all channels to have the noise-floor centered at 0 and the maximum level normalized to 1, to exclude the impact of relative brightness of each channel, prior to computing the cross-talk. We then used the scaled ground-truth images from the first 4 channels (leading channels) to produce ROIs around the beads using the *imbinarize* Matlab function, generating one set of ROIs for each leading channel. For each leading channel, we applied its ROIs to all trailing channels and summed the absolute values of these signal levels. Absolute values were used to also capture negative cross talk between channels in the cross-talk metric. We summed these numbers obtained from each leading channel for a given condition (ground-truth, de-mixed and raw data). For demixed and raw data condition this number is designed to contain all the additional signal in all channels that is generated due to cross-talk. It will also contain noise and signal from objects from trailing channels that are coincidentally located in the ROIs detected on the leading channels. We therefore computed the same metric on the ground-truth data to evaluate the expected level of this metric without cross-talk. In all three conditions data contains exactly the same sample and the same channels have been registered from raw or de-mixed condition to ground truth reference channels to have objects located in the same parts of the image.

#### Detection of frames corrupted by chewing on behavioral reward

Following a successful gap cross onto the reward position the animals were given a crispy froot loop as a reward. Chewing on the froot loop corrupted a subset of imaging frames beyond recovery, probably by generating high-frequency resonance in the MEMS scanner. The effected frames were <5% of the data and occurred exclusively following the reward delivery, while the animal was chewing. To detect the effected frames we calculated the frame-wise fast correlation (FFC). To calculate the FFC we first calculated the Pearson correlation between each frame and the template produced in the process of motion correction. Then, to remove the influence of slow image drift, we low-pass filtered the FFC trace (movmedian Matlab function with width of 100 frames) and subtracted this from the original FFC trace to generate the final FFC metric. We next fitted a two component Gaussian mixture model to the distribution of FFC values using the Matlab function fitgmdist. The component with the lower average FFC was manually confirmed to correspond well to the corrupted frames. Corrupted frames were then defined as all frames for which the FFC was lower than the lower component mean FFC plus two standard deviations of the lower component. When calculating neuronal or ROI fluorescence traces the corrupted frames were excluded with the trace interpolated between the neighboring non-corrupted frames.

#### Onset time of neurons within sequences of neuronal activity in PPC

To characterize the sequential activation of neuronal populations in PPC we used a triggered average of fluorescence trace of each neuron (segmented region of interest) on behavioral events. First the events were manually analyzed using the visible light cameras data during offline analysis. Crosses from all platforms to the floor or to other platforms, reward events and refusals were labelled. The imaging data was averaged in the period -20 s to +30 s around each class of behavioral events. For further analysis only data for the event “EAT” was presented in the current work. For a corresponding “RANDOM” data, the same number of events randomly chosen were used to generate the shuffled data as comparison. The first 10 seconds of each such average was used as baseline to assess the variability of the neuronal activity more than 10 seconds before the behavior. To do that we calculated average and standard deviation of the trace for each neuron then used these values to compute a Z-score for the resting 40 s of data (interval [-10, 30] around the detected behavior). We then applied a threshold on the data that had a Z-score above 3. The first such event with a duration larger than 0.5 s for for each neuron occurring in the given interval was then used as onset of the corresponding neuron within the sequence. All neurons without any detected events were not attributed any onset time and not considered for further sequence analysis. Raster plots with one averaged and binarized neuronal per row and ordered on the y-axis from lowest (top) to largest (bottom) onset time is shown in Fig. 5d and the corresponding shuffled version in Fig. 5e. The detected onset times for each neuron was labelled in red.

To build the cumulative probability function from all imaging sessions from all animals we combined the onsets of all sequence cells, as they were detected on the same time interval around the cross, then sorted them in ascending order and used these times as X-axis data, with the cumulative percentage on the Y-axis. The 95% confidence interval was produced using the Matlab function ecdf. The histogram data on Fig. 5g is the time-binned counts of the same pulled together from all imaging sessions onsets data. The error bars were computed assuming a locally uniform in time neuron onset density and therefore a Poisson distribution of the onsets for each given bin, giving the standard deviation of the distribution of square root of the mean.

To confirm the significance of the measured differences between behaviorally aligned trigger averages and the shuffled counterparts, for each considered time-bin (data was combined with a bin-width of 5 s) we computed the differences of neighboring onset times on the onsets data pulled from all imaging sessions. We had then a distribution of these differences for each time-bin for the behaviorally aligned data and for the shuffled data. We assessed the significance of the measured differences using the Wilcoxon rank sum test (Matlab function ranksum) between the behaviorally aligned distributions and the shuffled ones and statistical significance threshold was set to p-value < 0.05. The significantly different time-bins were marked with an asterisk above the histogram (Suppl. Fig. 6).

#### Threshold-based determination of activity and inactivity segments

To segregate the recorded neuronal signals into periods where neurons were active and inactive we adapted a previously described Ca^2+^-transient thresholding technique^56^. We first defined for each neuron or structure a baseline mean and standard deviation as mean and SD of the smallest 20 % of the ΔF/F0 data, and used these quantities to calculate a the data for each neuron or structure as a z-score. Periods of activity were then defined as the periods with a duration of at least 1 s for which the z-score exceeded 5 or was more negative than -5. Periods of inactivity were defined as the complement of the activity periods for each neuron or structure. For this procedure we smoothed the ΔF/F0 data using a moving average over a five frame window.

#### Analysis of influence of sustained 2PE and 3PE on indicators of neuronal health

Previous studies of the effects of sustained fluorescence imaging using multiphoton excitation sources have found that long term illumination is correlated with an increase in neuronal baseline fluorescence as well as an increase in the time constant of the decay of Ca^2+^-transients^41,42^.

To quantify changes in neuronal baseline (inactive) raw fluorescence indicative of neuronal damage that were independent of global changes in image brightness, we first calculated neuronal normalized raw fluorescence for each neuron. Normalized raw fluorescence was calculated for each neuron by first calculating global background fluorescence frame-wise as the mean raw fluorescence signal overall all pixels which were not contained within neuronal ROIs and subtracting this frame-wise from the neuronal ROI raw fluorescence average. We then calculated, for each neuron, the delta baseline (inactive) neuronal fluorescence as the difference between the mean normalized neuronal fluorescence for all segments defined as inactive in the last and first 20% of the recordings. For this analysis the raw data was smoothed using a moving average over a five frame window.

For the comparison of the decay time constants of Ca^2+^-transients we first identified transients with approximately comparable characteristics. The selection criteria for this analysis were 1) selected transients were isolated events with the previous preceding transient returning to baseline (inactive as defined above) at least 5 s prior to the onset of the transient of interest and the next following transient starting at least 5 s after the transient of interest ended, 2) amplitudes between 30 and 80 % ΔF/F0, 3) transient baseline fluorescence, defined here as the mean fluorescence in a 1s window prior to the start of the transient, between -20 and 20 % ΔF/F0 and 4) a transient rise time, the time from start to peak of the transient, of less than 2s. For the qualifying transients we next fit the decay phase of the transient, defined here as the data segment from the transient peak to the end of the transient (time at which the transient returned to inactive as defined above), with a single exponential of the form a·exp(-b·x) using MATLAB’s fit function. To ensure comparable and reasonable fits to the data we selected transients where the R-squared (coefficient of determination) for the fit was greater than 0.9, and for the qualifying transients calculated the decay time constant (τ) as -1/b. Finally, we compared the average τ for qualifying transients in the first and last 20 % of each recording for each neuron. For this analysis we smoothed the ΔF/F0 data using a moving average over a ten frame window.

## Acknowledgments

We thank Kay-Michael Voit for advice and discussion on data analysis. We thank Uwe Czubayko and Jeanine Klesing for expert technical help. We thank Ruth Pohle for assistance with editing. Funding from Max Planck Society. The funders had no role in study design, data collection and analysis, decision to publish or preparation of the manuscript.

## Author contributions

Experimental design AK, DJW, JNDK. Microscope hardware development AK, JS. Animal preparation DJW, CB, and AS. Data collection AK, DJW, CB and AS. Analysis design and implementation AK, DJW, JS and JNDK. Manuscript preparation AK, DJW, JS, and JNDK.

## Competing Interests

The technical developments resulting from this work have been filed as part of a patent application (PCT/EP2024/054016).

## Data availability

The data in the manuscript will be available in a Dryad repository at the time of publication.

## Supplementary Figures

**Supplementary Figure 1.**
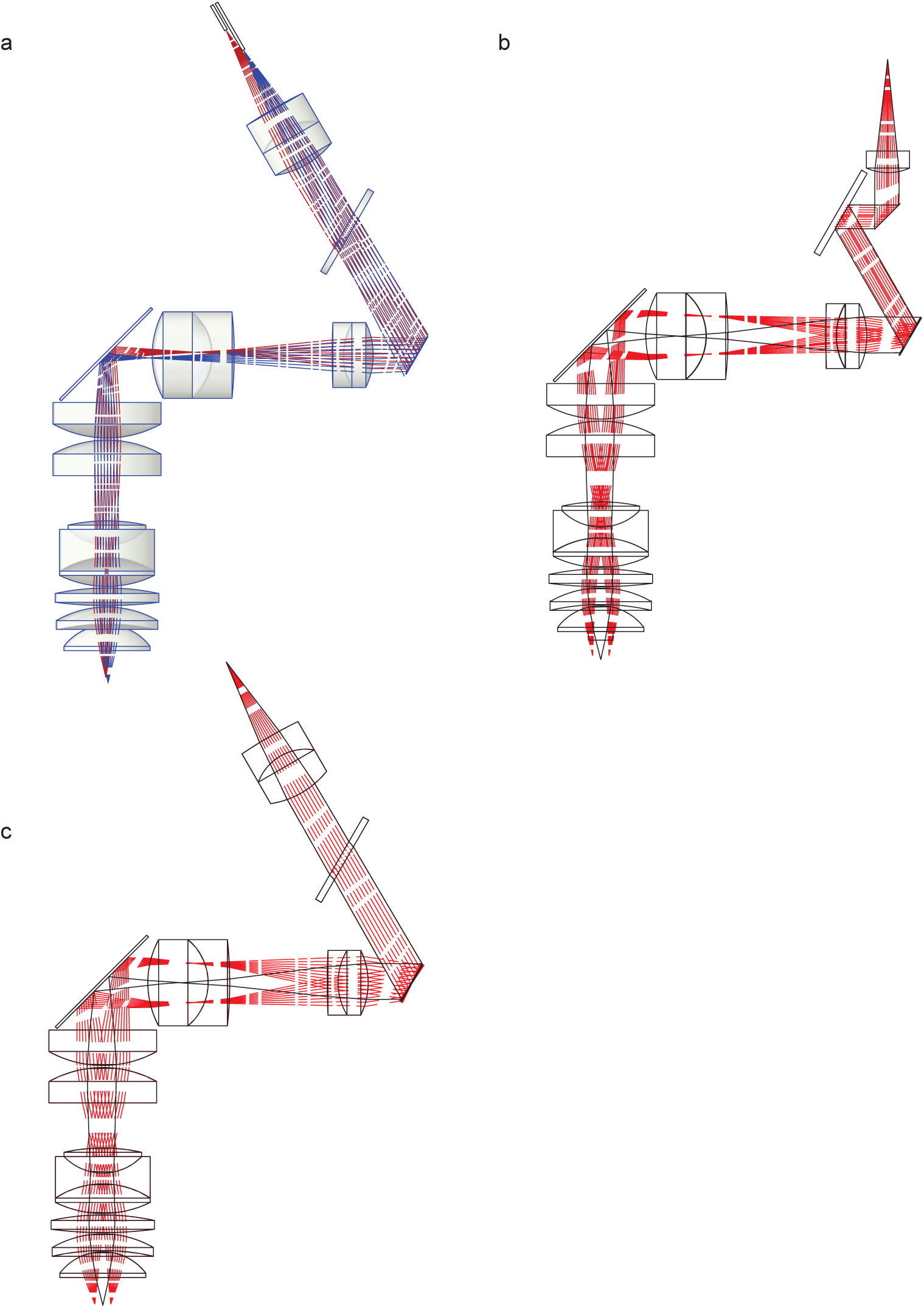
Zemax model for multimodality multiplane miniature microscope. **a**, scaled ray tracing from Zemax model featuring extreme positions of fibers to reach >250 µm axial offsets between imaging planes and corresponding lateral distance of 300 µm between fibers for unclipped beam propagation. **b**, scaled ray tracing from Zemax model with optical path for three-photon excitation, 1300 nm with MEMS scanner in the center position (black) and extreme field-of-view positions (red). **c**, scaled ray tracing from Zemax model with optical path for two-photon excitation, 960 nm with MEMS scanner in the center position (black) and extreme field-of-view positions (red).

**Supplementary Figure 2.**
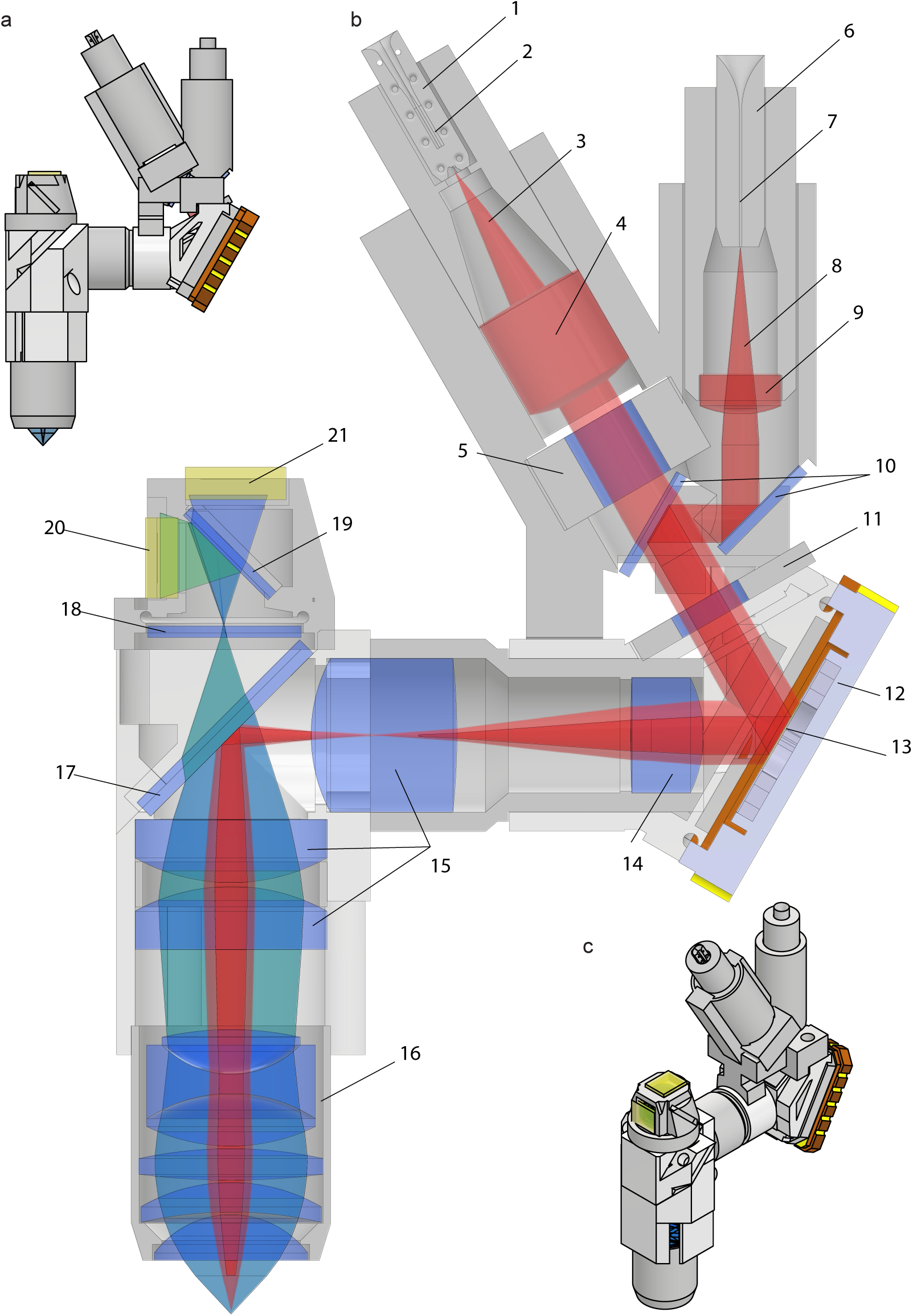
Optical and Electronic parts for assembly of the miniature multiplane microscope. **a**, side-view rendered from CAD microscope assembly. **b,** section-view of the miniature multiplane microscope along the symmetry plane with the detail of the components: 1, 3D printed ferrule for two-photon excitation (2PE) fibers, 2, individual bore for a 2PE fiber defining the 3D position of the fiber tip, 3, laser cone leaving the fiber tip of a 2PE fiber with the NA predicted from its core size at 960 nm, 4, achromatic doublet collimation lens, 5, 4-element stack ETL (µT Lens) for adjusting the axial offset between the three-photon excitation (3PE) plane and the group of 2PE planes, 6, ferrule for 3PE fiber, 7, fiber bore of the ferrule for 3PE, 8, laser cone leaving the fiber tip of the 3PE fiber with the NA predicted from its core size at 1300 nm, 9, aspheric collimation lens, 10, dichroic mirrors combining 2PE path and 3PE path, 11, 1-element ETL for fine tuning for same field-of-view retrieval, 12, MEMS package, 13, MEMS scanner mirror, 14, scan lens from previous work (Klioutchnikov 2022), 15, compound tube lens composed of an achromatic lens and two plano-convex lenses, 16, objective lens from previous work (Klioutchnikov 2022), 17, dichroic mirror separating excitation optical path and emission optical path, 18, infra-red emission filter, 19, dichroic mirror separating blue light from (third-harmonic generation) and green fluorescence from GCaMP7f, 20, SiPM for green channel, 21, SiPM for blue channel. **c**, oblique-view rendered from CAD microscope assembly.

**Supplementary Figure 3.**
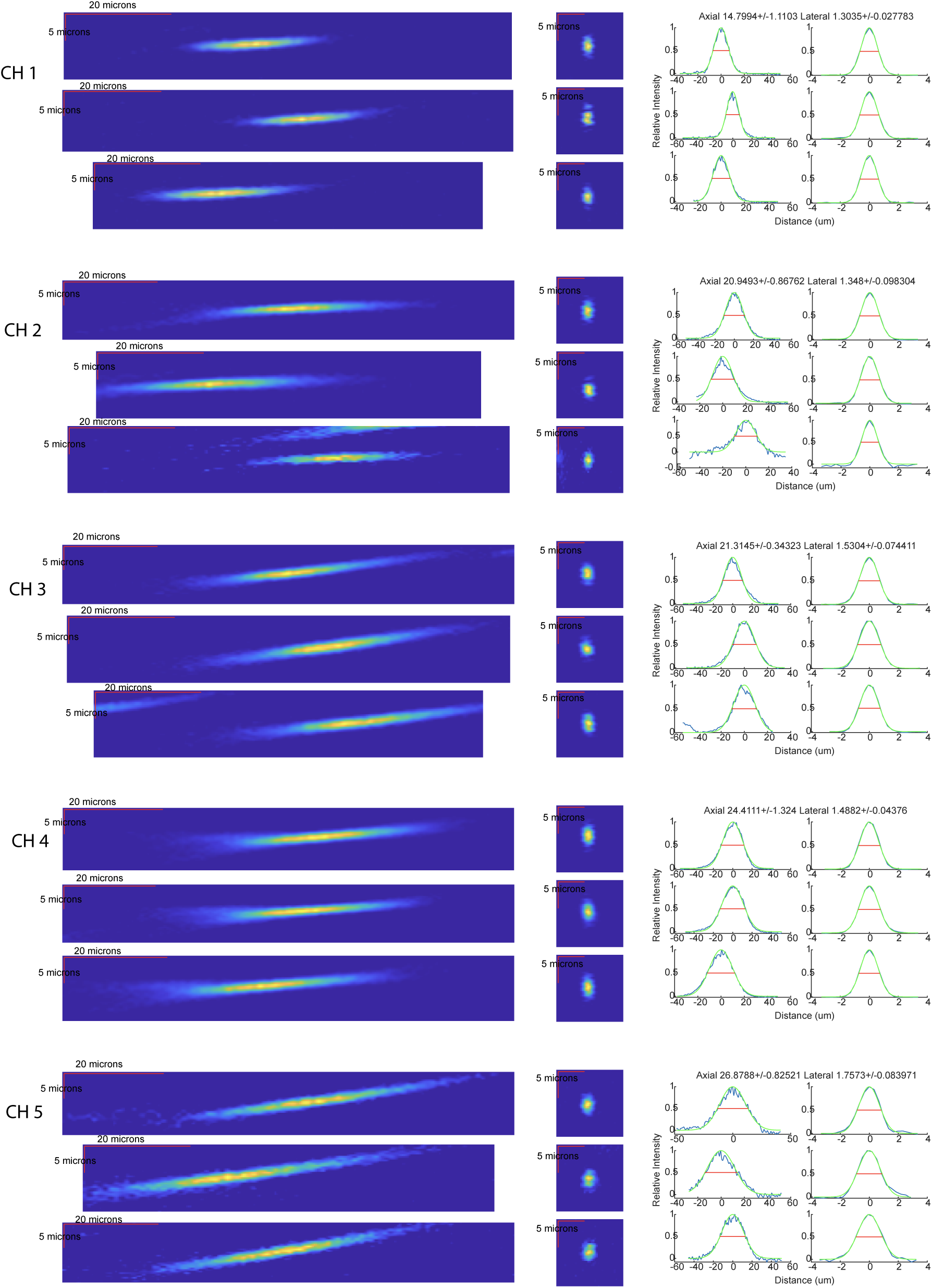
Optical resolution characterization. Point spread function measurement projection on a lateral dimension (slow axis, left column), the axial dimension (second column from left), projections on the axial dimension (blue) with Gaussian fit (green, second column from right) and projection on the fast axis (blue) and its Gaussian fit (green, rightmost column). All five channels have been separately characterized with the channel 1 on the top row and up to the channel 5 in the bottom row.

**Supplementary Figure 4.**
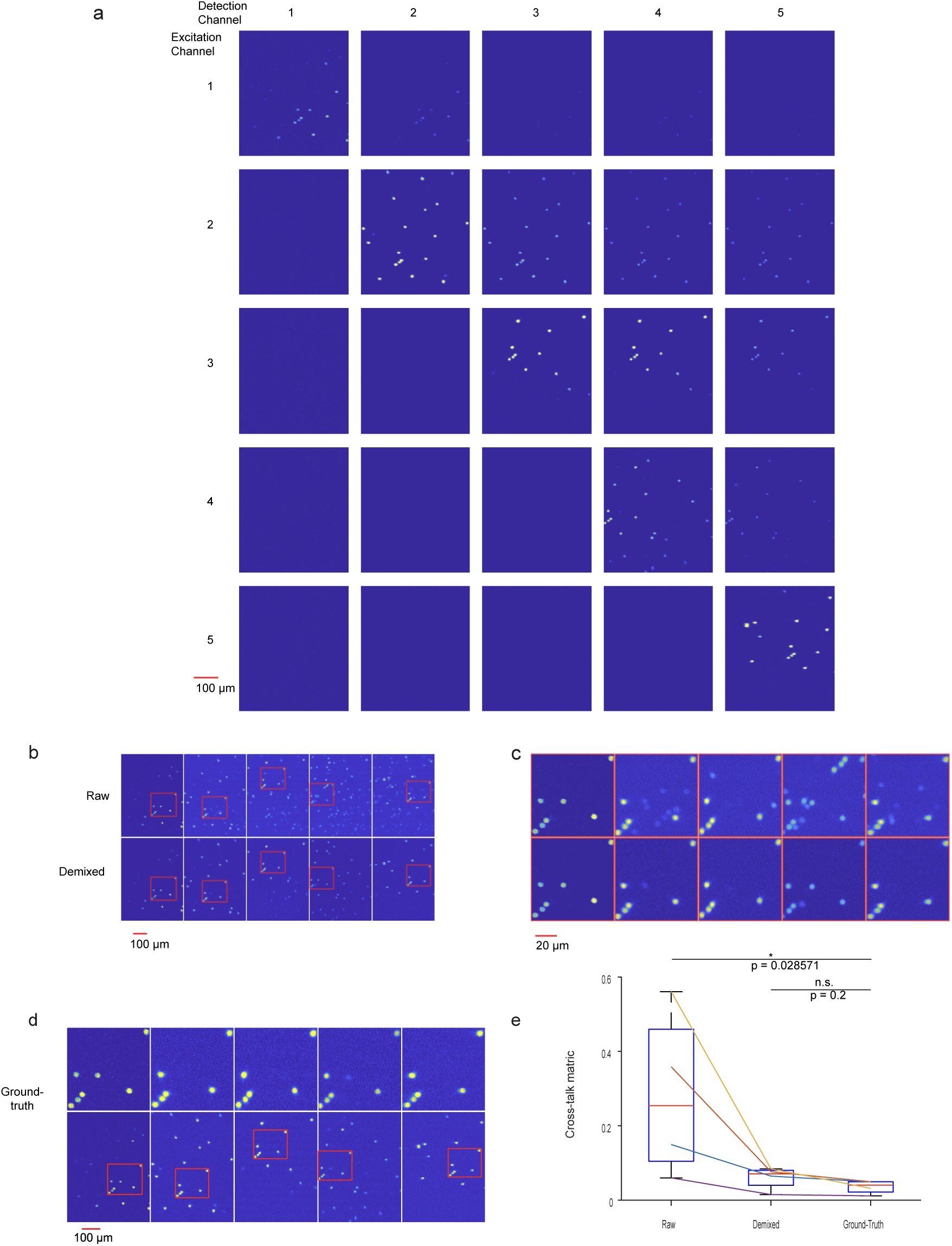
Ground truth data for de-mixing algorithm. **a**, summary of the de-mixing ground-truth data with the axial projections of the imaged 10 µm bead volumes with the only exciting fiber operational on the rows and the recorded channels on the columns. Note that recorded channels are ordered in the electronic order, such that each next channel is ∼ 8 ns later. On the diagonal we have the operational channel and the ground-truth signal and all the cross-talk is in the trailing channels, but never in the previous. **b,** axial projection of the same beads volume imaged as in a, but with all fibers operational. Raw data on the upper row and data after de-mixing algorithm in the bottom row. **c,** zoom on the same data as b, on the outlined in red areas, that were mapped in-between channel, where the same set of beads were found in each imaged volume. **d,** same axial projection from ground-truth data as in a, with the same beads outlined as in b and c and the corresponding zoom. **e,** cross-talk metric described in methods, that is the sum of absolute values of the signal in all detect ROIs for each channel, but computed in the trailing channels, to assess how much a given structure is producing signal in the trailing channels. Wilcoxon rank sum test was used to evaluate statistical significance.

**Supplementary Figure 5.**
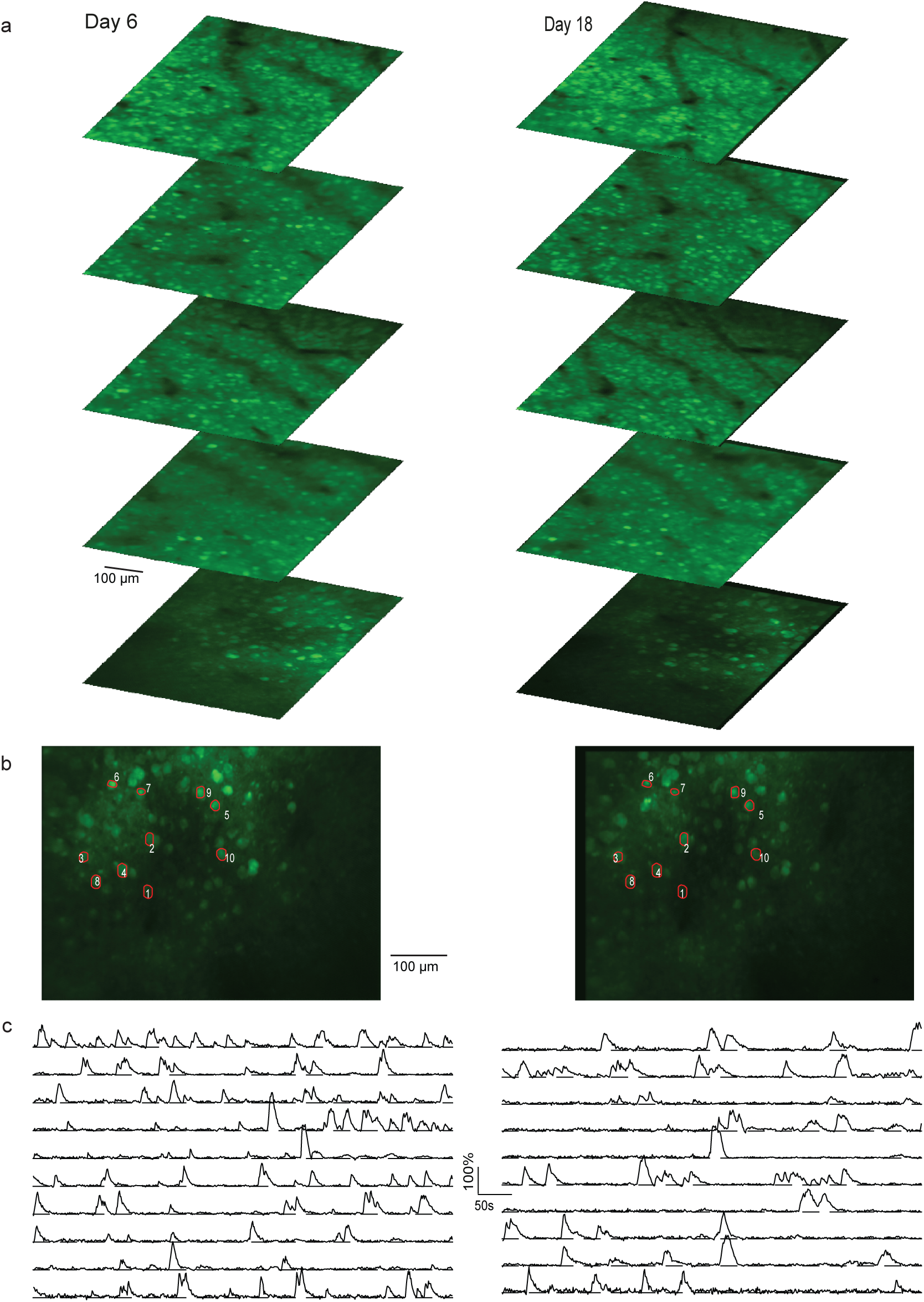
Multiplane imaging is stable over weeks. **a**, overview average images from all planes of labelled neurons in the posterior parietal cortex on day six (left) and near identical populations imaged on day 18 (right). **b**, overview images from channel five (3PE) in a, showing 10 example ROIs from which fluorescence traces are shown in c. **c** fluorescence traces corresponding to the ROIs labelled in b.

**Supplementary Figure 6.**
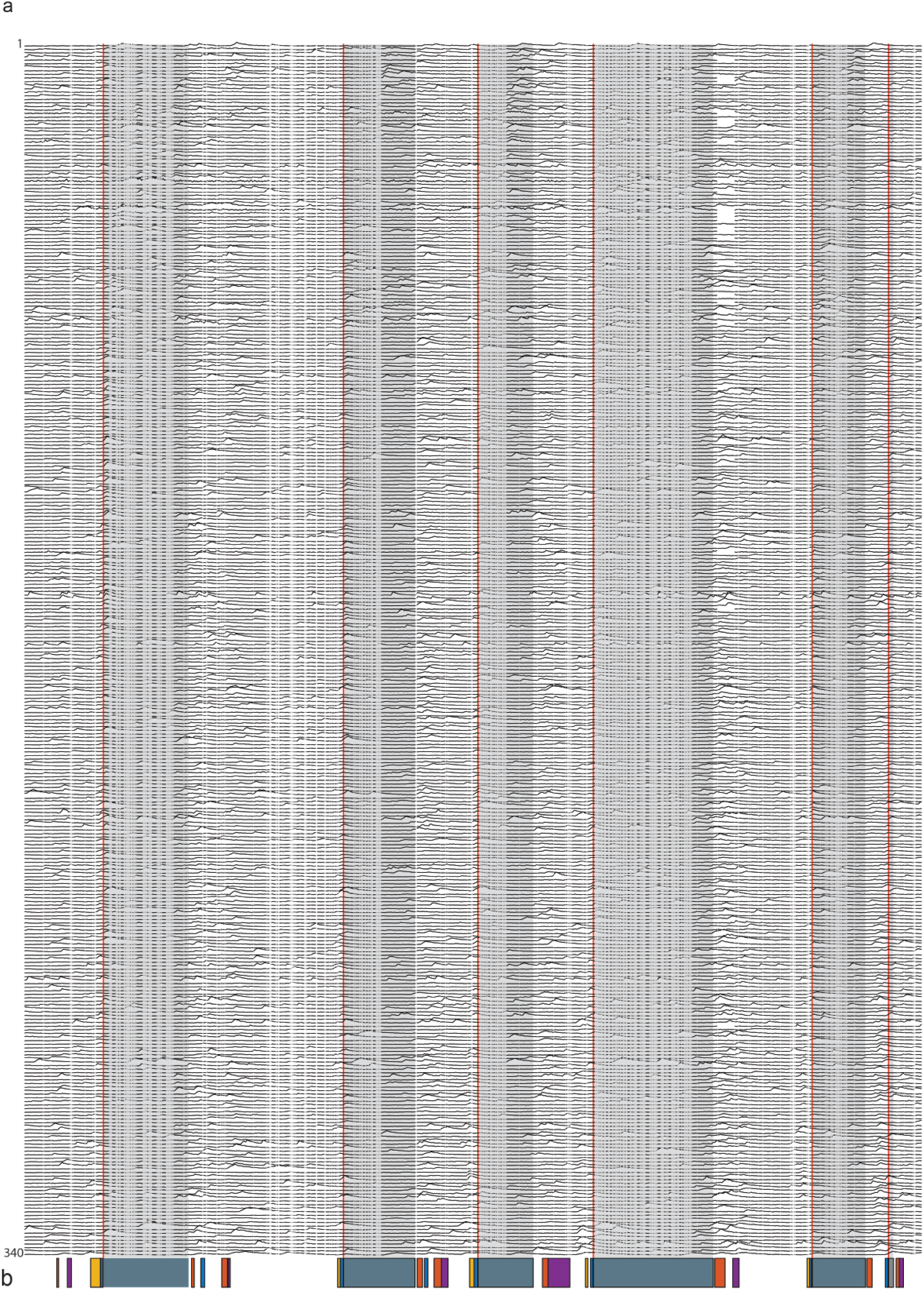
Activity of all sequence neurons from an imaging session. **a**, fluorescence traces of all sequence neurons detected in the imaging session described in Fig. 4 (N = 340 neurons). Red lines label gap-crosses. **b,** temporal representation of identified behavioral events in the recording session aligned on the same x-axis as a, (blue, cross to top platform, orange, cross to middle platform, magenta, cross from middle platform to arena floor, gray, reward location on top platform).

**Supplementary Figure 7.**
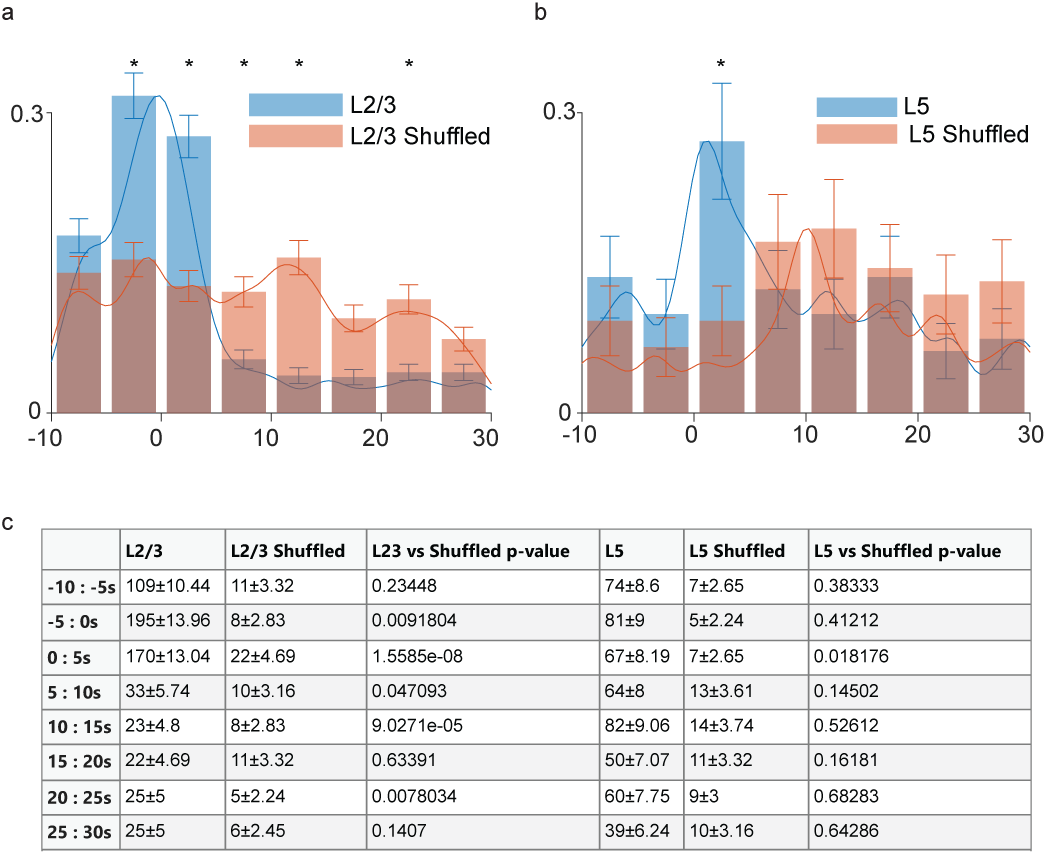
Statistical comparison between temporal density of sequence cells in L2/3 and L5 with shuffled data. **a**, probability density histogram of sequence cells in L2/3 and L2/3 Shuffled data, where exactly the same method was applied to detect sequence cells, on the same data with the difference that Shuffled data had the behavior event times randomly chosen trough the imaging sequence. Error bars show ±SD assuming Poisson distribution of sequence cells occurrences within each time-bin. Spline distribution fits shown as alternative continuous representation (ksdensity Matlab function was used to produce the fit on the histogram counts data). Asterisks mark statistical significance of the difference of distribution assessed in each time-bin (see Methods and c, for detail of statistical method). **b,** as in a but for sequence cells in L5 and L5 Shuffled data. **c,** summary of statistical data for histogram shown in a and b, for L2/3, L5 and their shuffled versions (format M±SD of the counts, each count being a detected sequence neuron, for probability density see a and b). To assess statistical significance, the difference between neighboring onset-times distribution was considered (inverse of distribution measures of instantaneous density). Wilcoxon rank sum test was used to assess significance of difference between medians of each bin of L2/3 and L5 data against the shuffled versions and each p-value is reported in a separate column. A threshold of 0.05 was set for statistical significance and marked by an asterisk in a and b.

